# SLC4A2 Anion Exchanger Promotes Tumor Cell Malignancy via Enhancing H^+^ Leak across Golgi Membranes

**DOI:** 10.1101/2021.02.09.428406

**Authors:** Elham Khosrowabadi, Antti Rivinoja, Maija Risteli, Anne Tuomisto, Tuula Salo, Markus J Mäkinen, Sakari Kellokumpu

**Author notes:** Corresponding authors: University of Oulu, Faculty of Biochemistry and Molecular Medicine, Aapistie 7A, PO BOX 5400, FIN-90014 Oulun Yliopisto, Oulu, Finland.

## Abstract

Proper functioning of each secretory and endocytic compartment relies on its unique pH micro-environment that is known to be dictated by the rates of V-ATPase-mediated H^+^ pumping and its leakage back to the cytoplasm via an elusive “H^+^ leak” pathway. Here, we show that this proton leak across Golgi membranes involves AE2a (SLC4A2a)-mediated bicarbonate-chloride exchange, as it is strictly dependent on both bicarbonate import (in exchange of chloride export) and the AE2a expression level in the cells. Imported bicarbonate anions and luminal protons then facilitate a common buffering reaction that yields carbon dioxide and water before their egress back to the cytoplasm via diffusion or water channels. The high surface-volume ratio of flattened Golgi cisternae helps this process, as their shape is optimal for water and gas exchange. Interestingly, this pathway is often upregulated in cancers and established cancer cell lines, and responsible for their markedly elevated Golgi resting pH and attenuated glycosylation potential. Accordingly, AE2 knockdown in SW-48 colorectal cancer cells was able to restore these two phenomena, and at the same time, to reverse cells’ invasive and anchorage-independent growth phenotype. These findings suggest that a malignant cell can be returned to a benign state by normalizing its Golgi resting pH.

## Introduction

Altered cellular metabolism, tumour acidosis and aberrant glycosylation are all hallmarks of cancers and contribute to tumorigenesis and its progression by various means [1–3]. Cancer-associated glycosylation changes most often include increased branching and/or fucosylation of N-linked glycans, synthesis of truncated mucin type O-glycans, increased levels of sialic acid as well as decreased sulphation of glycosaminoglycans such as heparan sulphate [4]. Because cell surface glycans regulate a vast number of different cell-cell and cell-matrix interactions [5,6], their alterations modulate a variety of basic cellular functions, including inflammatory responses, immune evasion, apoptosis, cell attachment as well as cancer cell dissemination, motility, invasion and metastasis [7–11].

Despite most cell surface glycans (collectively known as the glycocalyx) are made in the Golgi apparatus by the consecutive actions of Golgi-resident glycosyltransferases, the mechanistic details behind cancer-associated glycosylation changes are incompletely understood. Potential causes include altered expression of glycosyltransferases, their mislocalization, loss of catalytic activity, and inability to form functionally relevant complexes in the Golgi [12,13]. Moreover, previous work from our laboratory suggest that an abnormally high Golgi resting pH in cancer cells [14] appears to be one cause for altered glycosylation as well as protein sorting [15], given that pH gradient dissipating compounds such as chloroquine or ammonium chloride markedly increased the level of cancer-associated O-glycans in cells that normally express these glycotopes at low levels [14,16,17].

Golgi resting pH is normally acidic (pH 6.5-6.0) and set by three main ion transport systems: the vacuolar (V)-ATPase (that pumps protons to Golgi lumen), the GPHR chloride channel (Golgi pH regulator that prevents build-up of membrane potential brought about otherwise by proton pumping), and a proton leak “channel” that allows escape of protons back to the cytoplasm across Golgi membranes [18–21]. Of these, the proton leak pathway appears to be the most important one for setting the Golgi resting pH or “set point”, because its rate decreases along secretory pathway (ER, Golgi, secretory vesicles) concomitantly with their increasing acidity [22].

While both the V-ATPase and the GPHR protein are well characterized at the molecular level, the identity of the proton leakage pathway remains an enigma. Previous physiological measurements have suggested that proton efflux across Golgi membranes is voltage-sensitive and inhibited by Zn^2+^, suggesting the involvement of a regulated “channel”[19]. Other studies have proposed that proton leak involves NHE7 and NHE8 Na^+^/H^+^ exchangers [23]. However, a recent study shows that these exchangers act as acid loaders, and not as acid extruders, in the Golgi [24]. The AE2 (SLC4A2) bicarbonate-chloride exchanger isoform that is expressed in the Golgi membranes in a variety of cell types, could also contribute to Golgi resting pH by either importing or exporting HCO_3_^−^. The rationale behind this possibility is that all the four members of this Na^+^-independent Cl^−^-HCO_3_^−^ exchanger gene family (SLC4A1-4 or AE1-AE4) mediate a diisothio-cyanatostilbene-2,20-disulfonate (DIDS)-sensitive, electroneutral and obligatory one-to-one exchange of chloride for bicarbonate [25,26]. Thereby, they all regulate intracellular pH (pHi), cell volume, and chloride concentration in the cells.

The AE2 isoform is detected in nearly all tissues and cells examined, and therefore, is said to have a “house-keeping” role in the cells. Consistent with this, AE2 knockout in mice causes a severe phenotype, as AE2 (-/-) mice often die either before or at weaning [27]. AE2 has at least 3 known variant polypeptides (AE2a-c) which differ from each other by their variable N-terminal domains [26]. Both the N- and C-termini of the AE2 are known to be cytoplasmic and can interact with proteins that regulate either the activity or its subcellular localization in the cells. The C-terminal tail interacts with carbonic anhydrase II (or IV), an enzyme in the cytoplasm that produces bicarbonate anions (and protons) for transport [28–30], forming a “transport metabolon”. This interaction enhances AE anion transport activity as well as catalytic activity of the CAII [30]. The N-terminus of the AE2a (the longest N-terminal variant) in turn has been shown to bind ANK195, a Golgi-specific ankyrin isoform [31] that links the AE2-ANK195 complex to βIII spectrin-based Golgi membrane skeleton [32]. Thus, this interaction mediates AE2 localization in the Golgi.

To find out whether the AE2 isoform indeed contributes to Golgi resting pH in the cells, we utilized both AE2a overexpressing and knockdown cells as well as direct Golgi pH measurements in live cells using the ratiometric pH probe pHluorin [33]. We show that AE2a variant contributes to Golgi resting pH by enhancing proton leakage across Golgi membranes via the well-known bicarbonate-buffering reaction that yields carbon dioxide and water from imported bicarbonate anions and luminal protons. We also verified the functional relevance of this pathway by showing that this AE2-mediated pathway is often upregulated in cancers and in established cancer cell lines. Finally, we provide direct evidence to show that AE2 knockdown in SW-48 colorectal cancer cells is able to restore Golgi resting pH and glycosylation potential in the cells as well as to reverse their invasive and anchorage-independent growth phenotype. Glycan profiling of invasive and non-invasive SW-48 cells also revealed two potential glycotopes that may govern this phenotypic change between the cells.

## Results

### AE2a expression level alters Golgi resting pH and glycosylation potential

To investigate directly whether the AE2a variant regulates Golgi resting pH, we performed Golgi pH measurements in COS-7 cells using ratiometric, Golgi-targeted pHluorin as a probe. COS-7 cells express the Golgi-localized AE2a variant [31] and also display normally acidic (pH 6.3-6.5) Golgi resting pH [17] that is typical for mammalian cells [20]. Both immunofluorescence (Fig. 1A) and immunogold cryo-electron microscopy (Fig. S1A) with the AE2-specific antibody confirmed that AE2a its almost exclusive localized in the Golgi and especially in the medial- and trans-Golgi cisternae, and the trans-Golgi network. Ratiometric Golgi pH measurements in live COS-7 cells (GT-pHluorin, Fig. 1B) showed that forced overexpression of the AE2a-mCherry fusion protein (Fig. 1C) increased median Golgi resting pH by 0.5 pH units (pH 6.4 to pH 6.9) when compared to untreated or mock-transfected cells (Fig. 1D). Single-cell data analyses also revealed a strict correlation (R^2^ = 0.95) between the AE2a expression level and Golgi resting pH (Fig. 1E).

**Fig. 1.**
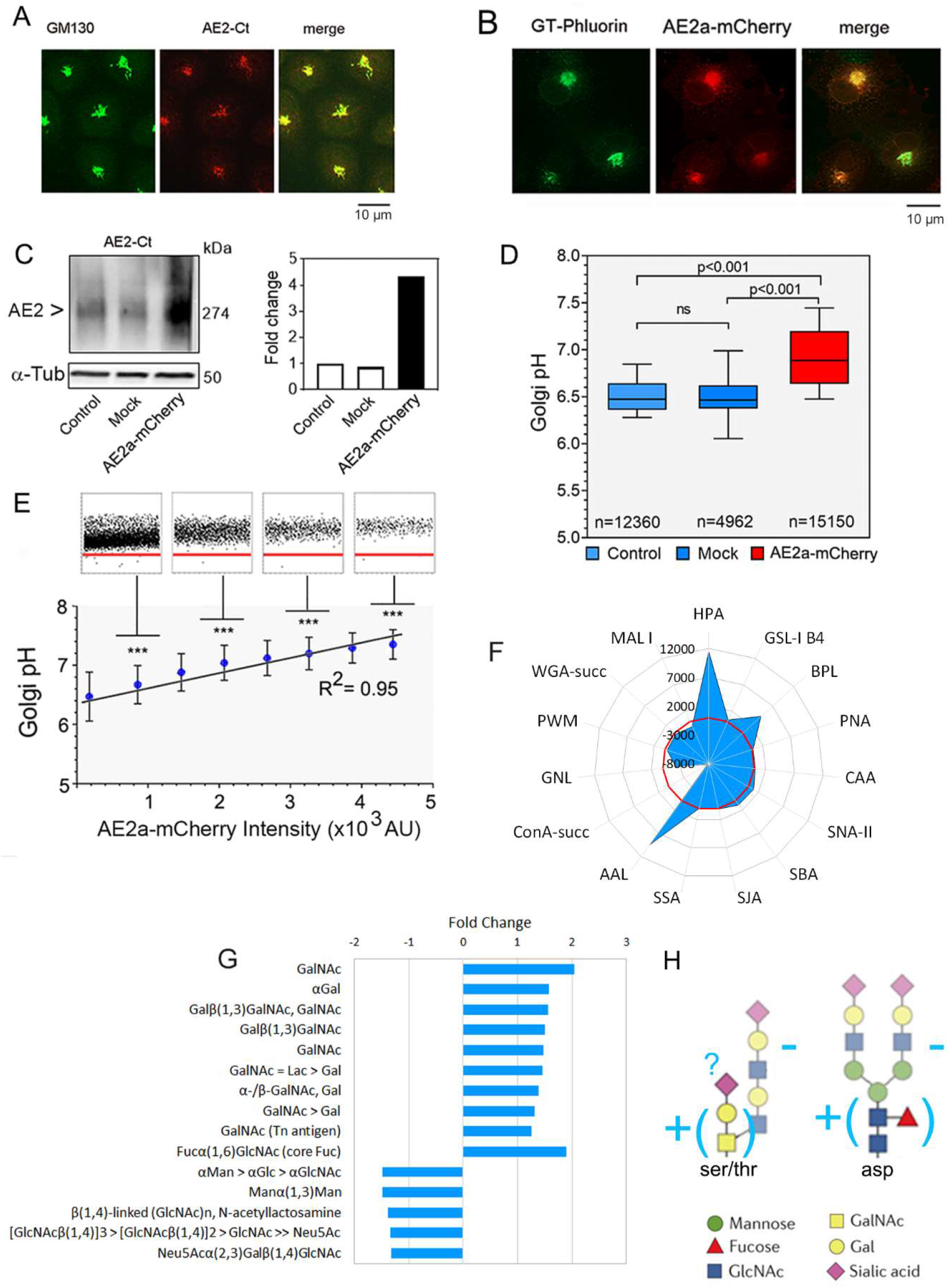
AE2a overexpression in COS-7 cells alters Golgi resting pH and its glycosylation potential. **(A)** Immunofluorescence microscopy of the endogenous AE2a protein and its co-localization with the Golgi marker (anti-GM130) in COS-7 cells. **(B)** Fluorescence microscopy of COS-7 cells transfected with the GT-pHluorin- and AE2a-mCherry-encoding plasmids. The merged figure shows their co-localization in the Golgi. (**C)** Quantification of the AE2a-mCherry protein level in wild type COS-7 cells (control), in mock-transfected cells and AE2a-mCherry expressing cells. The protein was visualized after Blue Native-PAGE (BN-PAGE) by immuno-blotting with the anti-AE2Ct antibody. (**D)** Ratiometric Golgi pH measurements in wild type, mock-transfected or AE2a-mCherry (red) expressing COS-7 cells. AE2a-mCherry overexpressing cells were selected using the Operetta TM build in software using the mCherry tag as a marker. In the box blot analysis, the whiskers indicate 10^th^ to 90^th^ percentiles; n = number of cells used for analyses. **(E)** Regression analysis between Golgi resting pH and AE2a-mCherry expression level. Single-cell data points (dots) were classified into eight equal classes based on AE2a-mCherry intensity (200-5000 AU units, 600 AU units/class), and plotted against the Golgi resting pH. Single cell data points are shown for every second class. The red line (pH 6.0) is added to help comparison. The means (± SD, n > 2000 cells) were used to obtain the fitted regression line and its coefficient (R^2^ = 0.95). **(F)** Lectin binding differences between AE2a-mCherry overexpressing COS-7 cells and mock-transfected control cells. Total binding intensities (Fig S1C) were used to calculate the means (± S.D, n=6) to select statistically significant (p < 0.05) changes between the cells. Two-tailed student’s T-test was used for statistical analysis. The means were then subtracted from each other (AE2a-mCherry minus control) to get the subtracted fingerprint shows as a radar plot. The red coloured ring denotes to zero line (no difference). (**G)** A bar graph showing significant differences (F) as fold changes. Lectin names were replaced with their sugar binding specificities. (**H)** A cartoon that illustrates main N- and O-glycan differences between AE2a-mCherry overexpressing cells and mock-transfected controls.

By using lectin microarray glycan profiling (Fig. S1B), we found significant (p < 0.05) differences with 15 out of 43 lectins between mock-transfected and AE2a-overexpressing COS-7 cells (Fig. 1F). The subtracted fingerprint revealed that several lectins that are specific for truncated O-linked glycan glycotopes such as GalNAc (or Tn-) and Galβ(1,3)GalNAc (or T-) glycotopes, were significantly upregulated in AE2a overexpressing cells, compared to mock-transfected controls (Fig. 1G). In addition, four lectins (Succinylated ConA, GNL, PWM, MAL I) that recognize terminal mannose-, N-acetylglucosamine-, galactose- and α(2,3)-linked sialic acid residues (Fig. 1G) showed that these N-glycan glycotopes were significantly decreased in AE2 overexpressing cells compared to mock-transfected controls. Only one lectin (AAL) that is specific for core-fucosylated N-glycans showed increased binding upon AE2a overexpression.

Collectively, these findings suggest that AE2a overexpression, besides increasing Golgi resting pH, also alters glycosylation either by increasing the synthesis of O-glycan core structures or by impairing their elongation (Fig. 1H). The fact that these glycotopes are normally masked by further glycosylation in the Golgi [11] supports the latter possibility. Altered N-glycans likely reflect their impaired processing and termination in the Golgi.

To provide further evidence for its role in Golgi pH regulation, we also knocked down the endogenous AE2a in COS-7 cells. To accomplish this, we employed doxycycline inducible SMART™ vector system, as it allows adjustment of the knockdown efficiency and thus, preservation of cell viability, in contrast to the observations with AE2 knockout mice [27]. In stably transfected and selected cells, each of the three AE2-specific shRNAs (Table S1) were able to decrease the level of the AE2a protein in COS-7 cells by 10-60% (Fig. S1C). Each one of them decreased Golgi resting pH by ~0.2 pH units upon induction compared to scrambled shRNA or the resting pH of non-induced cells (Fig. 2A). Single cell data analyses further confirmed that the AE2 knockdown affected uniformly all cells in each cell population (Fig. 2B). Lectin microarray analyses in AE2a knockdown cells (Fig. S1D) further uncovered significant (p < 0.05), yet less marked changes in glycosylation than that observed with AE2 overexpression. Nine (out of 43) lectins. All the lectins showed decreased binding intensities for a diverse set of glycotopes that range from the O-linked T-antigen, Blood group B glycotope to mannose-, N-acetylglucosamine- and α(2,3)sialic-terminating N-glycans (Fig.2C-D). These findings suggest that Golgi glycosylation potential is strictly dependent on its resting pH and that it needs to be strictly regulated around pH 6.3 to sustain efficient processing and elongation of both O- and N-linked glycans.

**Fig. 2.**
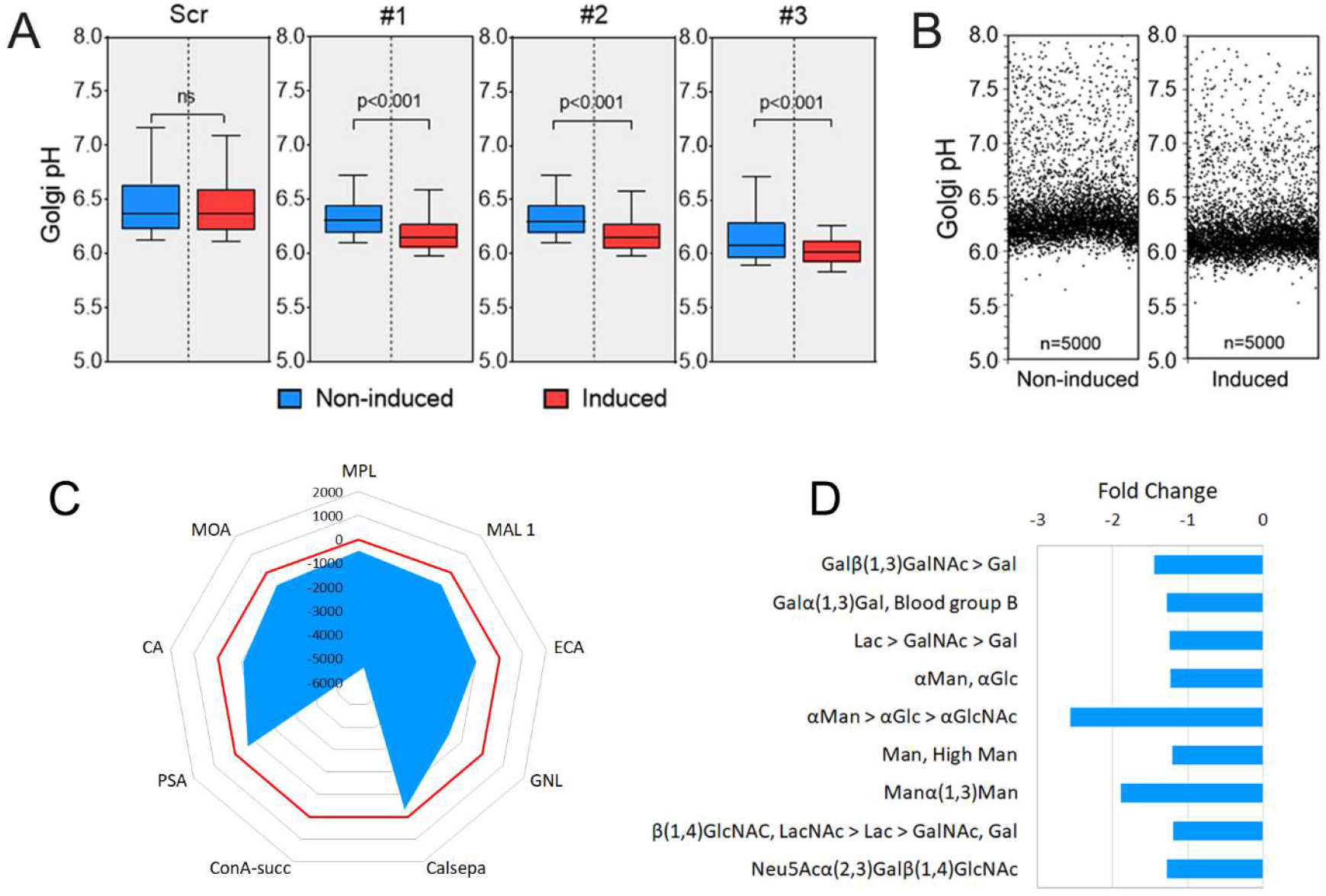
AE2a knockdown decreases Golgi resting pH and alters glycosylation. **(A)** Golgi resting pH in AE2a knockdown cells. Stably AE2 shRNA expressing COS-7 cells were transfected with the GT-pHluorin plasmid, and either induced (red boxes) or not (blue boxes) before determining their Golgi resting pH by ratiometric imaging (n= 5000 cells/ group). The whiskers above and below the box indicate 10^th^ to 90^th^ percentiles. (**B)** Single-cell analysis of the COS-7 cells stably expressing the shRNA#1 plasmid. The Golgi resting pH of both non-induced and induced cells is shown. **(C)** Subtracted fingerprint showing significant (p < 0.05) glycosylation differences between AE2a knockdown (induced) and control (non-induced) cells. Significant differences were calculated as above and are show as a radar plot. The red circle denotes to zero values (no difference). **(D)** A bar graph showing significant differences (C) as fold changes. Sugar specificities correspond to lectin names shown in (C) starting from MOA lectin to the left.

### AE2a enhances proton leakage rate across Golgi membranes

To assess why AE2a overexpression increases Golgi resting pH, or knockdown decreases it, we performed next Golgi acidification and proton-leakage rate measurements in COS-7 cells that were transfected either with an empty plasmid (control), with the AE2a-mCherry construct (AE2plus) and AE2a knockdown (KD) cells. To accomplish this, each group of cells was transfected with the ratiometric Golgi targeted GT-pHluorin before permeabilizing their plasma membrane with streptolysin O (SLO) to allow controlled addition of ATP, Cl^−^ or HCO_3_^−^ anions while leaving Golgi membranes intact [34]. Ratio-imaging of the cells were first done in Cl^−^ /HCO_3_^−^ free bath medium to minimize AE2a anion exchange activity. Addition of excess ATP (10 mM) to bath medium resulted in significant acidification of the Golgi lumen (Fig. 3A top panel), plateauing at pH 5.0-5.5 in control and AE2a overexpressing cells. The corresponding initial acidification rates (ΔpH/min) were 1.0 and 1.9 pH units/min, respectively (Fig. 3A bottom panel). Proton leakage rates measured after blocking the V-ATPase activity with Concanamycin A (CMA, 1 μM) were low (Fig. 3A middle panel), initial leakage rates ranging between 0.2-0.35 pH units/min (Fig. 3A, middle and bottom panels).

**Fig. 3.**
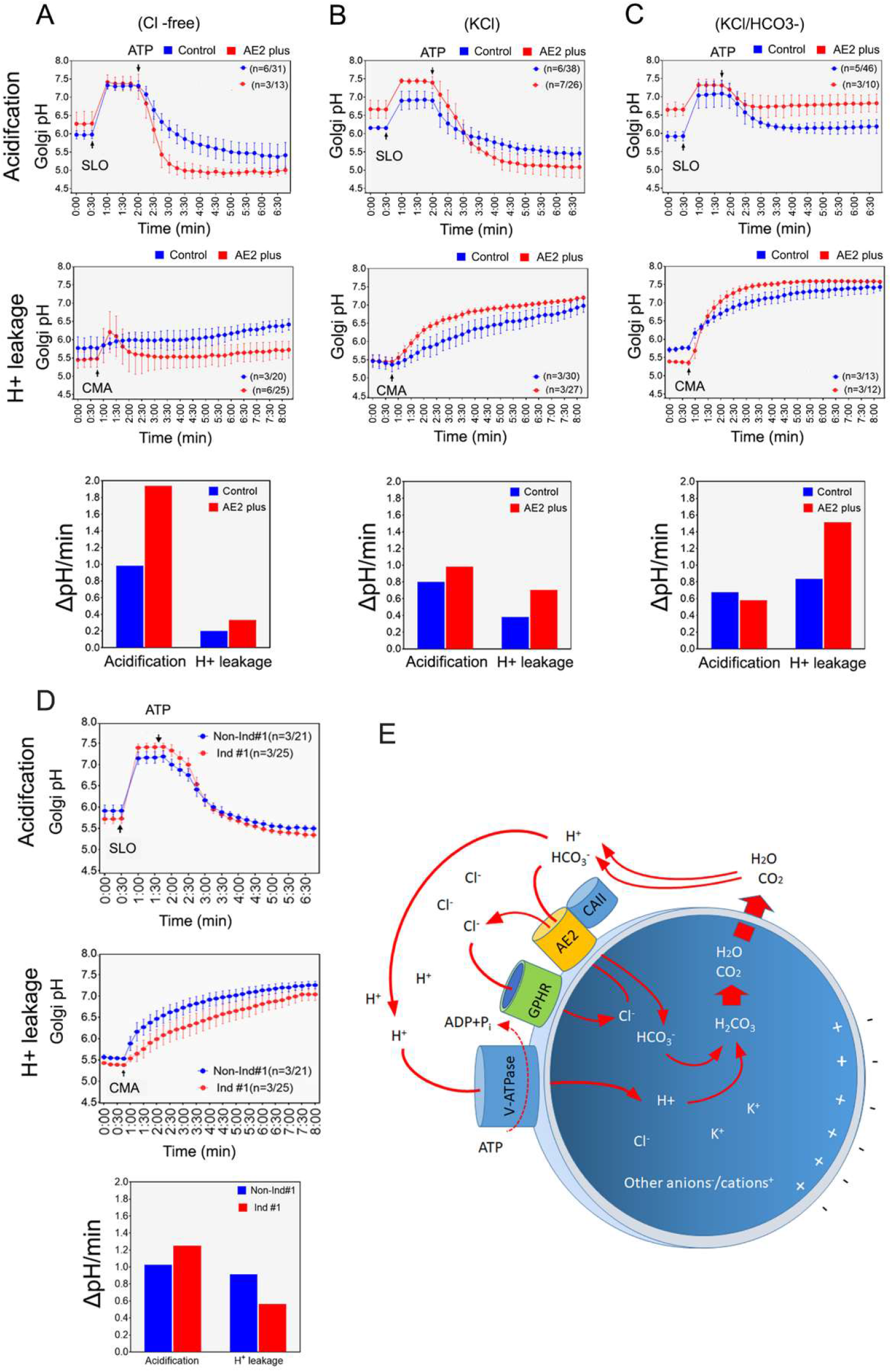
AE2a alters Golgi resting pH via facilitating proton leakage across Golgi membranes. Cells transfected with GT-pHluorin plasmid, either with or without the AE2a-mCherry plasmid, were permeabilized with SLO before ratio-imaging of the cells. Cells were kept in different bath solutions during the experiment: **(A)** no Cl^−^ nor HCO_3_^−^, **(B)** only Cl^−^ (no HCO_3_^−^) and **(C)** both Cl^−^ and HCO_3_^−^. Arrows denote the addition time of SLO, ATP or CMA to the bath medium. The obtained ratios (mean ± SD, n ≥3, marked in each graph) were transformed to pH values using the calibration standard curve and its equation before plotting against time. The bars (bottom panels) depict the initial acidification and leakage rates (ΔpH/min) that were calculated from the fitted curves in each case. (**D)** Golgi acidification and proton leakage rates in AE2 knockdown (non-induced) COS-7 cells and in control (non-induced) cells carrying the same plasmid (shRNA#1). The pH values were determined as above and plotted against time. The initial Golgi acidification and proton leakage rates (ΔpH/min) were also determined as above. **(E)** A schematic cartoon representing AE2a-mediated proton leakage pathway across the Golgi membranes. The main ions as well as corresponding transporters are shown. AE2 and CA II form the transport metabolon via interacting with each other.

Addition of Cl^−^ anions to the bath medium (by replacing KCl with K-gluconate) had no marked effect on the initial acidification rate in wild type cells (1.0 units/min vs 0.8 units/min, (Fig. 3B top and bottom panels) while in AE2a overexpressing cells, ΔpH/min was decreased from 1.9 ΔpH/min (no chloride) to 1.2 ΔpH/min (with chloride, Fig. 3B bottom panel). Proton leakage rates were increased in both control and AE2a overexpressing cells by approximately 2-fold compared to those in chloride-free bath medium (Fig. 3B, middle and bottom panels). This increase probably is due to the presence of endogenous HCO_3_^−^(produced likely by the CA II) as AE2 cannot transport chloride without bicarbonate [26]. As expected, AE2a overexpressing cells showed higher H^+^ leakage rate than mock-transfected control cells (Fig. 3B, middle panel).

When both Cl^−^ and HCO_3_^−^(20 mM) were present in the bath medium (Fig. 3C), Golgi acidification rates in both control and AE2a overexpressing cells were decreased, plateauing at pH 6.3 (control cells) and at pH 6.7 (AE2a overexpressing cells). These changes coincided with their markedly increased proton leak rates (Fig. 3C middle and bottom panels), AE2a overexpressing cells displaying nearly 2-fold higher leak rate than mock-transfected control cells. These findings suggest that proton leakage rate across Golgi membranes is strictly dependent on the presence of both bicarbonate and chloride anions as well as the abundance of the AE2 protein in the Golgi membranes. In support for this view, AE2a knockdown cells displayed significantly lower proton leakage rate (Fig. 3D) and slightly higher acidification rate than non-induced control cells (Fig. 3D). We also excluded the possibility that the above changes are caused by bicarbonate-mediated changes in cytoplasmic pH, as none of the pre-calibrated HCO_3_^−^-free buffer solutions (pH 6, pH 7, and pH 8) added to SLO-permeabilized cells did not affect Golgi resting pH (Fig. S2A). On the other hand, when we cultivated intact COS-7 and HeLa cells (Fig. S2B and C) in the presence of excess bicarbonate (40 mM), we noticed a small but significant (p<0.001) increase in Golgi resting pH in both cell types.

Collectively, these findings suggest a model (Fig. 3E and supplementary video) in which AE2a-mediated bicarbonate import (in exchange for luminal chloride) facilitates proton leakage across Golgi membranes by enhancing the formation of carbon dioxide and water from luminal bicarbonate anions and protons. This reaction (H_2_O + CO_2_ < > H_2_CO_3_ < > H^+^ + HCO_3_^−^), also known as the bicarbonate-buffering reaction [35] is directed towards left due to continuous AE2-mediated bicarbonate import and also by the egress of water and carbon dioxide across Golgi membranes. This can occur either via diffusion or aquaporin water channels, as the latter is able to transport both H_2_O and CO_2_ [36,37]. This process is likely helped also by the flattened shape of the Golgi cisternae, as their high surface-to-volume ratio (similar to erythrocytes) can markedly accelerate water and gas transport across lipid membranes. It is perhaps not a mere coincident that the pKa of the bicarbonate buffering system (pKa 6.1) is in same range with the Golgi resting pH (pH 6.5-6.0).

### AE2 protein is often upregulated in cancers and in established cancer cell lines

To assess whether the AE2a-mediated proton leakage pathway described above has any functional relevance in cancer cells that display elevated Golgi resting pH [17], we screened first Cancer Genome Atlas (TCGA) mRNA expression database using the GEPIA web server [38]. Since the mRNA analyses cannot discriminate between the different AE2 variants (AE2a-c), a term “AE2” will be used from here onwards. We found that the AE2 mRNA levels are significantly (p < 0.05) upregulated in 16 out of the 31 (52%) different cancer types present in the database. Of these, six cancer types showed highly significant (p < 0.001) AE2 upregulation (Fig. 4A). Kaplan-Meier survival plots of these six cancer types also suggested that high AE2 mRNA expression level is associated with markedly shorter life span of cancer patients (Fig. 4B). Pan-cancer analysis with all 31 different cancer types (Fig. 4C) gave similar, yet less marked results. Patient survival time was shortened from 95 months to 70 months.

**Fig. 4.**
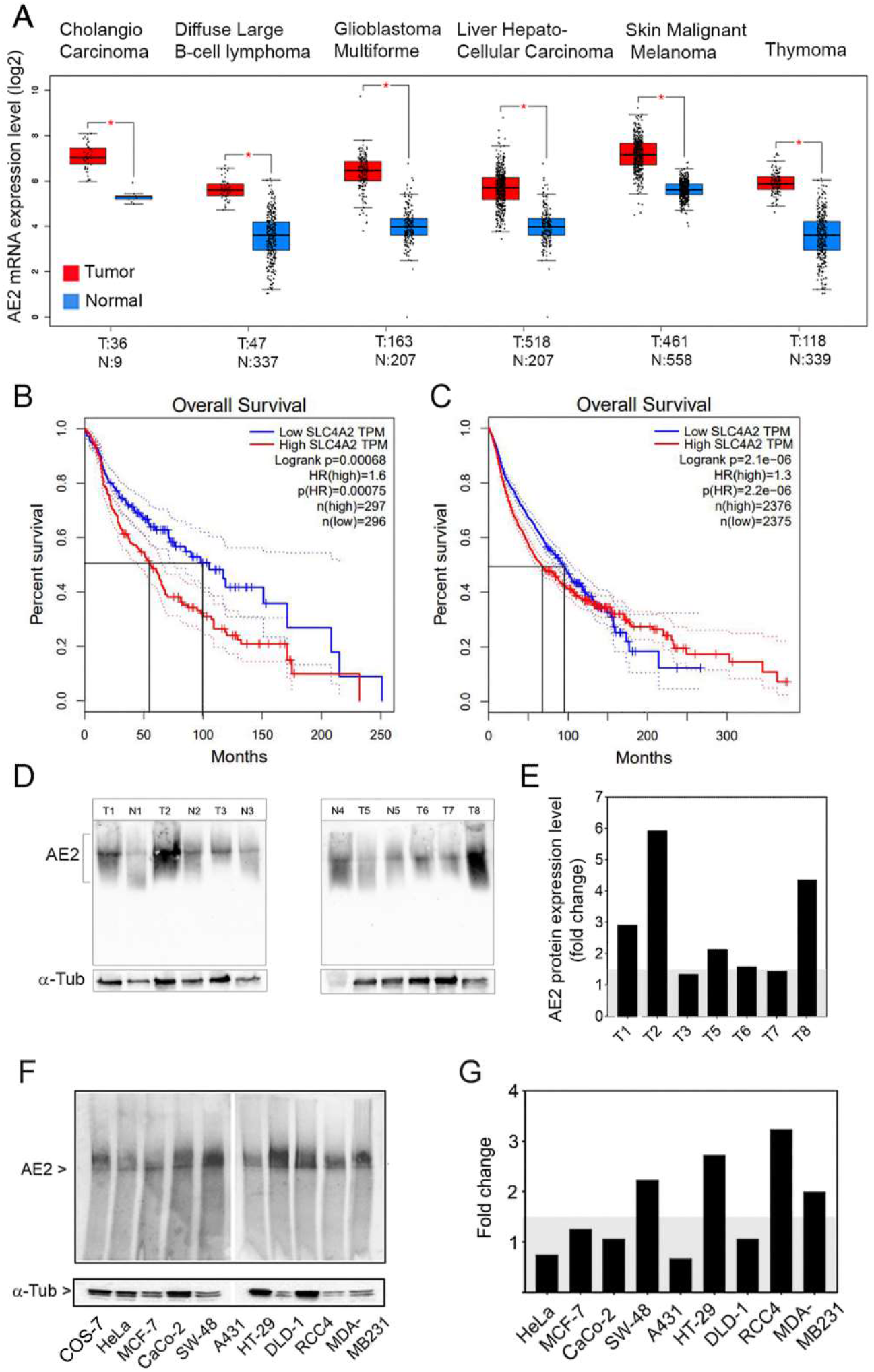
AE2 mRNA and protein expression levels in human cancers and established cancer cell lines. (**A**) The Cancer Genome Atlas (TCGA) and the GTEx databases were analysed for AE2 mRNA expression level using the GEPIA web server interface (38) and pre-set settings (jitter size 0.4, log2FC cut off 1, p-value cut off 0.01). The figure shows six specific cancer types (out of 31) that display highly significant (p < 0.001*) up-regulation of the AE2 mRNA relative to normal tissue samples from the same patient. (**B)** Kaplan-Meier overall survival plots of the above six cancer types. **(C)** Kaplan-Meier survival plot from all 31 cancer types (pan-cancer analysis). In both B and C, 75% and 25% cut-offs were used for high and low expressors, respectively. (**D-E**) AE2 protein expression levels in seven distinct colon cancer patient tissue specimens. All cancer tissue specimens with low-grade histology (as well as their healthy control tissue from the same patient) were evaluated by a clinical pathologist for grading. Immunoblotting with the anti-AE2Ct antibody was used for quantification of the AE2 protein level relative to healthy tissue sample except in patient samples T7 and T8 for which healthy tissue controls were not available. Hence, the mean of the other five normal patient samples were used for comparison. ImageJ was used for the quantification of the protein band intensities from the blots. All intensity values were normalized against α-tubulin. **(F-G)** AE2 expression levels in different cancer cell lines. Protein levels were determined and quantified as above after immunoblotting with the anti-AE2Ct antibody.

Further support for the AE2 upregulation in cancers was obtained also at the protein level. Immunoblotting after Blue-Native PAGE of colorectal cancer tissue samples with low grade histology using an affinity-purified anti-AE2 C-terminal antibody revealed that AE2 expression level varies between the patients by 5-fold when compared with each other (Fig. 4D, E). Comparison each cancer tissues sample to the corresponding healthy tissue showed that in four out of seven (57%) cancer specimens, the AE2 protein expression level was more than 1.5-fold higher than in normal tissue samples. Similar results were obtained by immunoblotting of total cell lysates prepared from nine different cancer cell lines (Fig. 4F, G). Overall, the AE2 expression level varied 6-fold between different cancer cell lines. When compared to COS-7 cells (which display normal Golgi pH, Fig. 1D), four cancer cell lines (44%) displayed more than 1.5-fold higher AE2 protein level than did COS-7 cells (Fig. 4F-G).

### AE2 overexpression is responsible for elevated Golgi resting pH in SW-48 cancer cells

The observed upregulation of the AE2 protein in about half of the cancer tissue or cell samples tested, lead us next to examine whether it has any consequences on Golgi resting pH. Golgi pH measurements performed as above uncovered that Golgi resting pH was markedly elevated in all 9 different cancer cell lines (Fig 5A) while the reference cells (COS-7) display normal Golgi resting pH. Interestingly, Golgi resting pH values were highest (close to or above pH 7) in the four cancer cell lines (SW-48, HT-29, RCC4 and MDA-MB231) which also displayed highest AE2 protein levels (Fig. 4G). In the other cancer cell types, Golgi resting pH ranged between pH 6.4 (HeLa) and pH 6.8 (A431). Single-cell data blots (Fig. S3B) further demonstrated that the high variation seen especially in HeLa and MCF-7 cells is due to the presence of two distinct cell populations with different Golgi resting pH values. Their presence was not associated with cell death, as dead cells were typically lost during the measurements due to repeated washings and media changes, nor with fragmented Golgi elements typical for cancer cells (Fig. S3A), as both compact and fragmented Golgi elements had rather similar mean Golgi resting pH values (Fig. S3B). Moreover, nearly all HeLa cells display a compact Golgi morphology (Fig. S3A and S3C). Instead, their co-existence likely reflects adaptation to hypoxia, as exposure of MCF-7 and SW-48 cells to hypoxia (5% O_2_ instead of 16 % O_2_) for 48 h decreased Golgi resting pH only in cells which display Golgi resting pH higher than pH 7 (Fig. S3D). Hypoxia exposure, however, did not decrease Golgi resting pH of the main cell population.

**Fig. 5.**
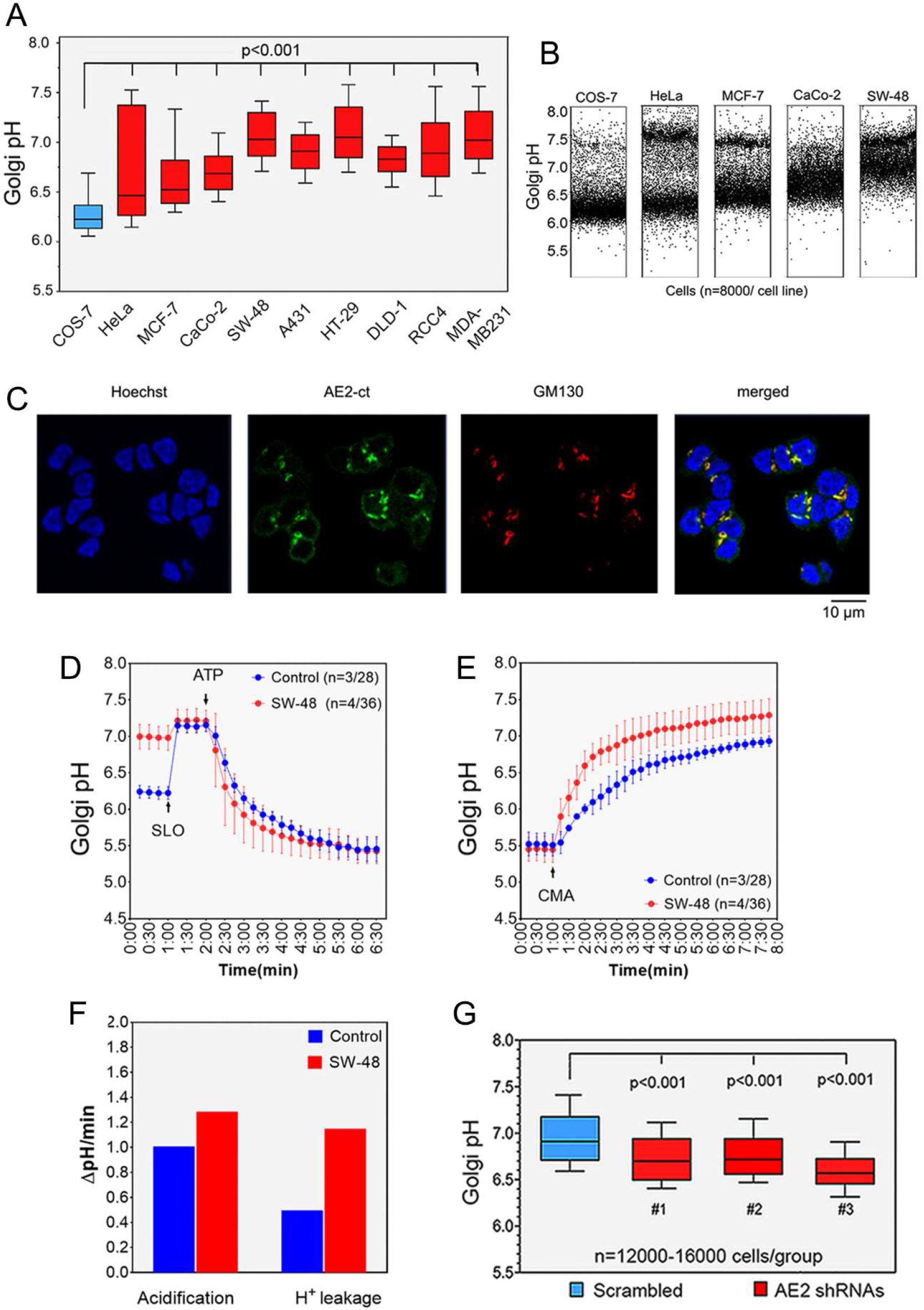
AE2 upregulation in cancer cells is associated with markedly elevated Golgi resting pH. (**A)** Golgi resting pH in different cancer cells types in comparison to non-malignant cells. Golgi resting pH in each cell type was determined as above (Fig. 1) using the GT-pHluorin as a probe. The values are shown using box plot graphs. Whiskers above and below each box indicate 10^th^ to 90^th^ percentiles. (**B)** Single cell Golgi pH data with four depicted cancer cell types. Golgi pH values from 8000 cells were blotted individually for each cell type. In three cell types, cells form two distinct cell populations with differing pH values. **(C)** Immunofluorescence microscopy of the endogenous AE2 protein showing its localization in the Golgi of SW-48 cells. Fixed cells were co-stained with the anti-AE2Ct antibody and with a Golgi marker antibody (GM130). **(D-F)** Golgi acidification (D) and proton leakage (E) rates in SW-48 cells. The experimental set-up is identical to that described in Fig. 3. The calculated initial acidification and leakage rates (ΔpH/min) are also shown (F). **(G)** Golgi resting pH in SW-48 cells after AE2 knockdown. Stably transfected SW-48 cells carrying one AE2-specific shRNA (#1-3) each or the control plasmid (scrambled shRNA) were induced for 3 days using 100 ng/ml of Doxycycline. Golgi resting pH values were then determined as above (e.g. Fig 1). The whiskers above and below the box indicate 10^th^ to 90^th^ percentiles.

SW-48 colorectal cancer cells were chosen for further analyses, mainly because 98 % of the cells display markedly elevated Golgi resting pH and AE2 protein (Fig. S3E). Most cells (75%) also stain positively with PNA lectin (Fig. S3F), a sign of impaired glycosylation (16). The mean Golgi resting pH of the two distinct cell populations was in both cases close to neutral, i.e. pH 6.9 and pH 7.4 (Fig. 5B). Our ATP level measurements also excluded the possibility that cells don’t have enough ATP to sustain proton pumping (Figs. S3G-H).

To verify that the upregulated AE2 in SW-48 cells is indeed responsible for their elevated Golgi resting pH, we first stained cells with the AE2Ct antibody and found that the AE2 protein is localized almost exclusively in the Golgi membranes of the cells (Fig. 5C). Subsequent Golgi acidification and proton leakage measurements revealed that upregulated AE2 protein did not alter Golgi acidification rate markedly (Fig. 5D). In contrast, the initial proton leakage rate (Fig. 5E-F) across Golgi membranes was 2.6-fold higher in SW-48 cells than in COS-7 cells (ΔpH 1.15/min vs. ΔpH 0.45/min, respectively). In addition, AE2 knockdown with the three AE2-specific shRNAs was found to decrease both theAE2 protein level in the cells (Fig. S4) as well as their median Golgi resting pH (Fig. 5G). Doxycycline induction with the shRNA#3 was the most effective one. It decreased median Golgi resting pH by 0.4 pH units (pH 6.5) compared to scrambled shRNA construct (pH 6.9).

Collectively, these data suggest that that enhanced proton leak due to AE2 upregulation is responsible for the elevated Golgi resting pH in SW-48 cells. In further support of this view, we did not detect any marked decrease in proton pumping rates between the cells (Fig. 5D) which suggest that the V-ATPase is fully functional in SW-48 cells.

### AE2 upregulation in SW-48 cells controls their invasive and anchorage-independent growth phenotype

To test whether AE2 upregulation has any consequences on cancer cell behaviour, we next performed cell proliferation rate measurements in wild type SW-48 (WT), in AE2 knockdown cells (KD) and in cells stably transfected with the scrambled shRNA construct (Scr). AE2 knockdown had no marked effect on cell proliferation rate when compared with either WT or Scr cells (Fig. 6A). Based on a scratch wound healing assay, AE2 decreased cell migration rate by 2-fold (Fig. 6B-C). AE2 knockdown cells also grew in a more dispersed manner relative to each other and did not anymore form cell clumps typically seen control SW-48 cells (Fig. 6D).

**Fig. 6.**
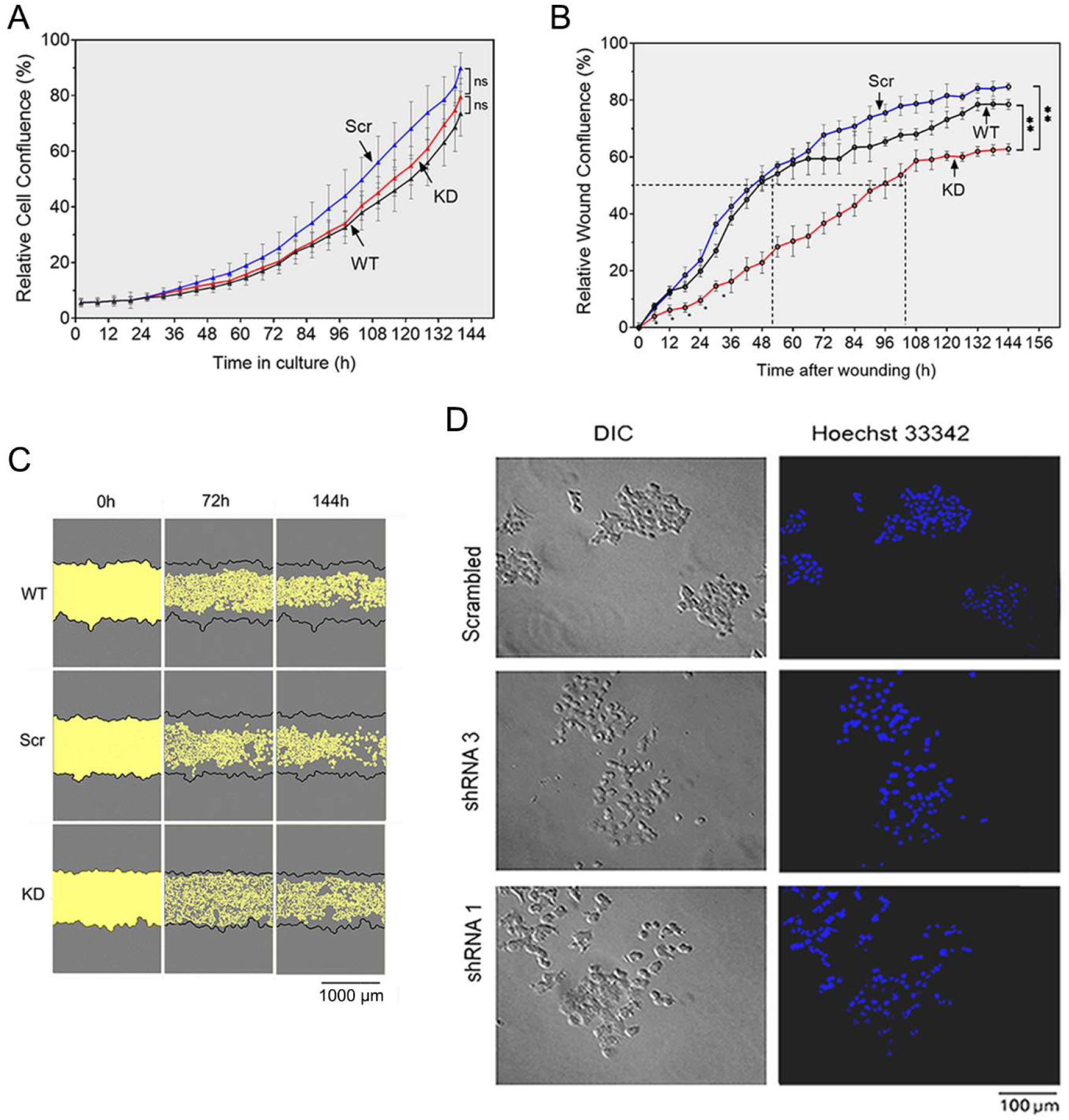
AE2 knockdown modulates SW-48 cell mobility and growth phenotype without affecting proliferation. **(A)** Cell proliferation rate measurements in AE2 knockdown cells in comparison to wild type SW-48 cells and knockout controls (SW-48 Scr). IncuCyte® live cell imaging system was used for the experiments. The means (± S.D, n = 9) of each time point were calculated using instrument’s own software package before plotting against time. For AE2 knockdown, cells carrying the shRNA#3 were used. **(B)** Cell motility rate measurements using the scratch wound healing assay. Confluent wells were scratched with an automated pin system, after which wound healing rate was followed by measuring cell confluence in the wound area. Cell confluency at each time point is shown as percentages of the wound area. **(C)** Representative figures on cell confluency are shown for each cell type. Cell-free areas are pin-pointed by yellow colour. (**D**) Growth phenotype of AE2 knockdown cells carrying either the shRNA#1 or the shRNA#3 construct in comparison to control cells (SW-48 WT and SW-48 Src). In brief, cells were induced for 3 days with doxycycline, then fixed and stained with Hoechst 33342 DNA dye before imaging. Representative figures are shown for each cell type. Differential interference contrast (DIC) microscopy was used for imaging.

Because SW-48 cells are known to be tumorigenic (https://www.lgcstandards-atcc.org/products/all/CCL-231.aspx#characteristics), we next measured their invasive potential by using an established 3D human myoma tissue based invasion assay [39]. Wild type SW-48 cells showed invasion foci inside the myoma tissue at 200 to 400 μm distance from the seeded top cell layer (Fig. 7A). Similar foci were also seen with SW-48 Scr cells (Scr) with or without doxycycline induction (Fig. 7B, C). In contrast, in AE2 knockdown cells, invasion foci were detected before the induction, but not after induction, suggesting that upregulated AE2 in SW-48 has a role in promoting SW-48 cell invasiveness. We also tested the opposite, i.e. whether non-invasive COS-7 cells become invasive upon AE2 overexpression. COS-7 cells stably expressing the AE2a variant in the Golgi displayed invasion foci while in mock-transfected cells, no invasion foci were detected (Fig. 7D, E). These findings suggest that AE2 upregulation in SW-48 cells is crucial for their high invasive potential.

**Fig. 7.**
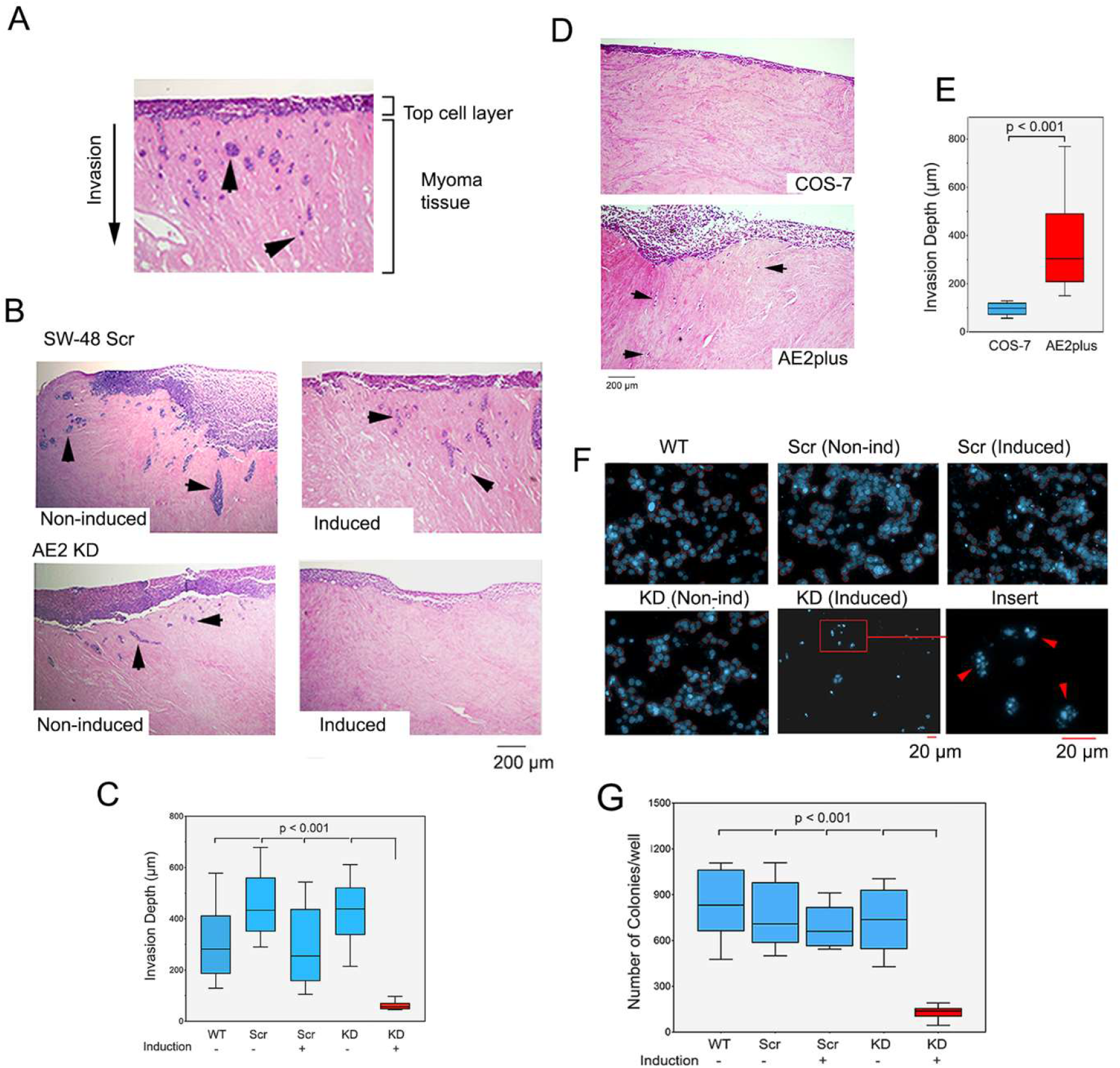
AE2 knockdown reverses the invasive and anchorage-independent phenotype in SW-48 cells. **(A)** Myoma-tissue based 3D invasion assay. Cells were seeded on top of myoma discs, allowed to grow for 21 days before fixing and processing for immunocytochemistry. Sections were then cut perpendicularly to the seeded cell layer, stained and imaged for quantification. A representative figure is shown. Invasive foci that are detected inside the myoma tissue (arrow heads) represent invaded cells. **(B)** Invasive potential of AE2 knockdown cells (KD) and control cells (SW-48Scr) were measured as above either with (Induced) or without (Non-Ind) doxycycline induction. Invasion depth was used as a measure of invasive potential. **(C)** A box plot presentation showing the median depth of the invasion foci from the top cell layer. Twelve sections from each myoma disc (n=2/24) were used for the quantification with ImageJ software. The whiskers indicate 10^th^ to 90^th^ percentiles. **(D)** Invasive potential of wild type COS-7 cells and AE2 overexpressing COS-7 cells (AE2plus). The invasion foci consisted mostly of single cells of which few are marked by arrows to pinpoint their existence. Note their absence in wild type COS-7 cells. **(E)** Median invasion depth in COS-7 cells shown as a box plot presentation. The whiskers indicate 10^th^ to 90^th^ percentiles. **(F)** Colony formation assay in soft agar. Depicted cells (wild type SW-48, WT; AE knockdown SW-48 cells, KD; SW-48 control cell, Scr) were grown in soft agar for 30 days with or without doxycycline induction (KD and Scr), after which they were fixed and stained with Hoechst dye before imaging with the Operetta high content screening system. The number of cell colonies present were quantified using the Harmony software. More than three nuclei in the pre-set area size was used to depict one colony. The insert (bottom, right) shows a higher magnification of the AE2 knockdown cells with fragmented nuclei (arrowheads). **(G)** Total number of colonies is shown as a box plot presentation where the whiskers represent 10^th^ and 90^th^ percentiles.

Another characteristic feature of a cancer cell is its ability to grow and proliferate without attachment to the substratum. Therefore, we employed a colony formation assay in soft agar to test whether AE2 upregulation also contributes to this phenotype. The data (Fig. 7F, G) showed that unlike control cells (SW-48 WT cells or non-induced and induced SW-48 Scr cells), AE2 knockdown cells were able to form colonies only in the absence of doxycycline, but not in its presence. In addition, AE2 knockout cells were present at low numbers, and also displayed fragmented nuclei (Fig. 7G, insert), suggesting that the remaining cells represent apoptotic cells. This may reflect loss of anoikis-resistance in AE2 knockdown cells, a special survival program that allows SW-48 cells to proliferate without any surface contact. Collectively, these findings support an important role for AE2 in the control of SW-48 cell malignant behaviour.

### AE2 upregulation enhances the synthesis of tumour-associated glycans in SW-48 cells

To uncover why AE2 knockdown reversed the SW-48 cell invasive and anchorage-independent growth phenotypes, we focused on potential glycosylation changes that can distinguish non-invasive cells (AE2 KD, COS-7) from the invasive ones (SW-48, SW-48 Scr). The rationale behind this is that cancer-associated glycosylation changes are well known to promote cancer cell invasiveness and metastatic spread [9–11, 40–42]. Moreover, glycosylation potential of the Golgi is highly dependent on Golgi resting pH [13,14,16,17] which is elevated in invasive SW-48 WT and SW-48 Scr cells, while the pH is normal in non-invasive AE2 knockdown cells and COS-7 cells (Fig. 5A and 5G).

The overall glycosylation patterns were rather similar despite some observable changes between the cells (Fig. S5A-B). Heat map analyses (Fig. 8A) showed that the highest and lowest binding intensities were detected with the same lectins listed on top and bottom of the map, respectively (Fig. 8A). To find out the main differences between non-invasive and invasive cells, we compared AE2 knockdown cells (KD) to control SW-48 Scr cells. Similar comparison was done also between wild type SW-48 and COS-7 cells. The main differences as shown as subtracted fingerprints for each pair (Fig. 8B). Statistical analyses further showed that out of the 43 lectins in the array, 24 showed statistically significant (p < 0.05) changes between AE2 knockdown (KD) and SW-48 Scr control cells (Fig. 8C). Similar analyses with wild type cells (SW-48 WT, COS-7) revealed differences in 17 out of 43 lectins (Fig. 8D). Of these, 9 lectins were common to both cell pairs (Fig. 8E) and could thus be regarded as important for cancer cell invasiveness. Further comparison of the two cell pairs with other revealed that only two lectins (ACA and RCA 120) behave identically in each pair. ACA lectin, which is specific for the T-antigen was markedly decreased, while RCA 120 which recognizes galactosylated complex type N-glycans was increased. Thus, these two lectins can distinguish non-invasive from invasive SW-48 cells.

**Fig. 8.**
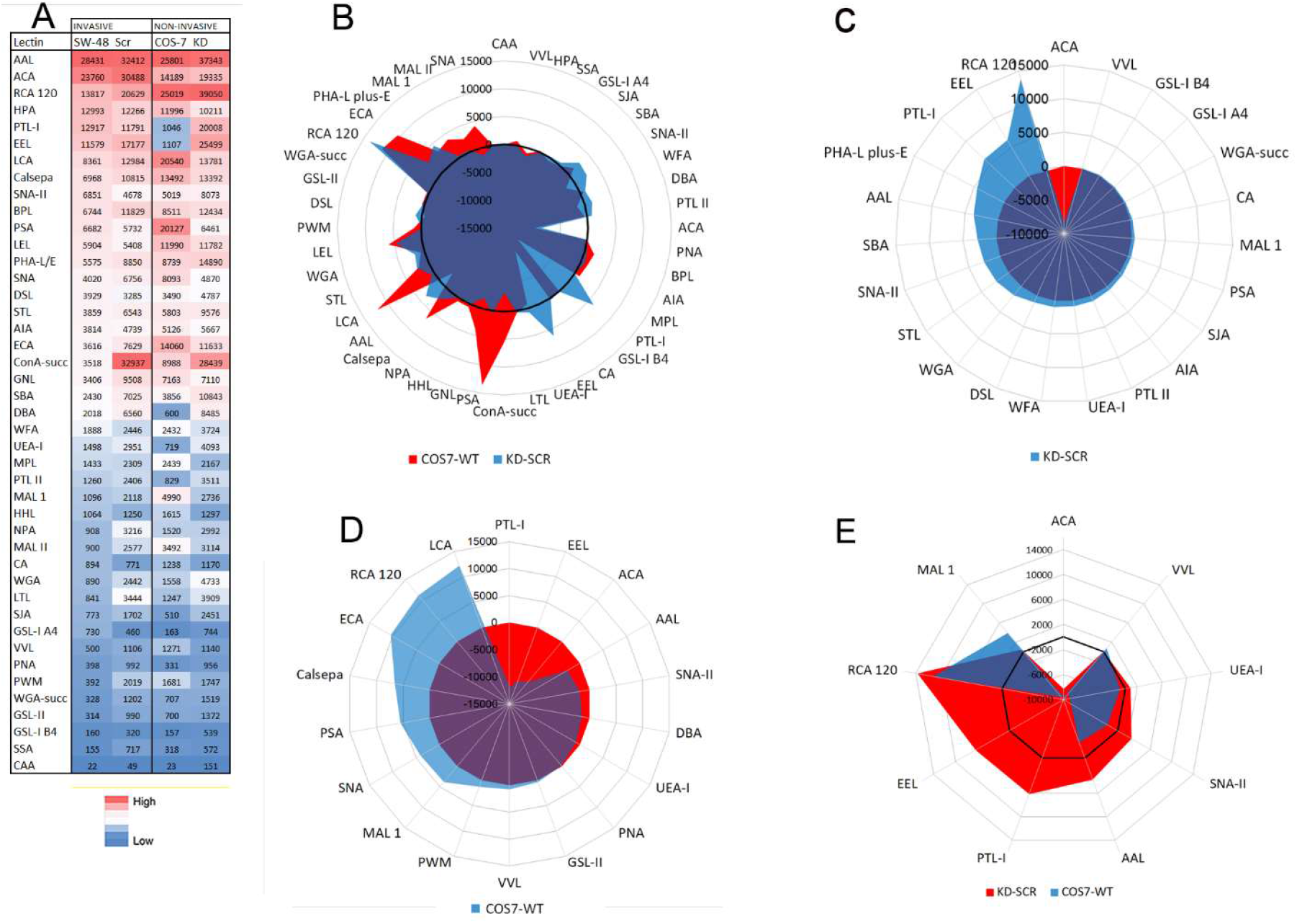
Glycan profiles differ between non-invasive and invasive cells. **(A)** Heat map analysis showing total lectin binding intensities after classifying them from largest to smallest using wild type SW-48 cells as a reference. For each cell type, a heat map was constructed separately using the conditional formatting in Excel. The means (Fig. S5) were used for calculations. **(B)** Glycan binding differences between COS-7 and SW-48 WT cells or between AE2 knockdown cells and control cells (KD vs. SW-48 Scr). Subtracted fingerprints are shown as a radar plot. The black-coloured circle denotes to zero values (no difference). **(C)** A radar plot showing statistically significant (p < 0.05) glycan binding differences between AE2 knockdown cells and control cells. Red colour in the radar plot depicts a decrease and light blue an increase. **(D)** A radar plot showing a subtracted fingerprint of statistically significant (p < 0.05) changes between COS-7 cells and wild type SW-48 cells. The same colour codes were used as above. **(E)** Comparison of subtracted fingerprints between non-invasive and invasive cell type pairs (AE2 knockdown cells vs. SW-48 Scr and SW-48 WT vs. COS-7). Only common changes and the corresponding lectins are shown on the radar plot.

**Fig. 9.**
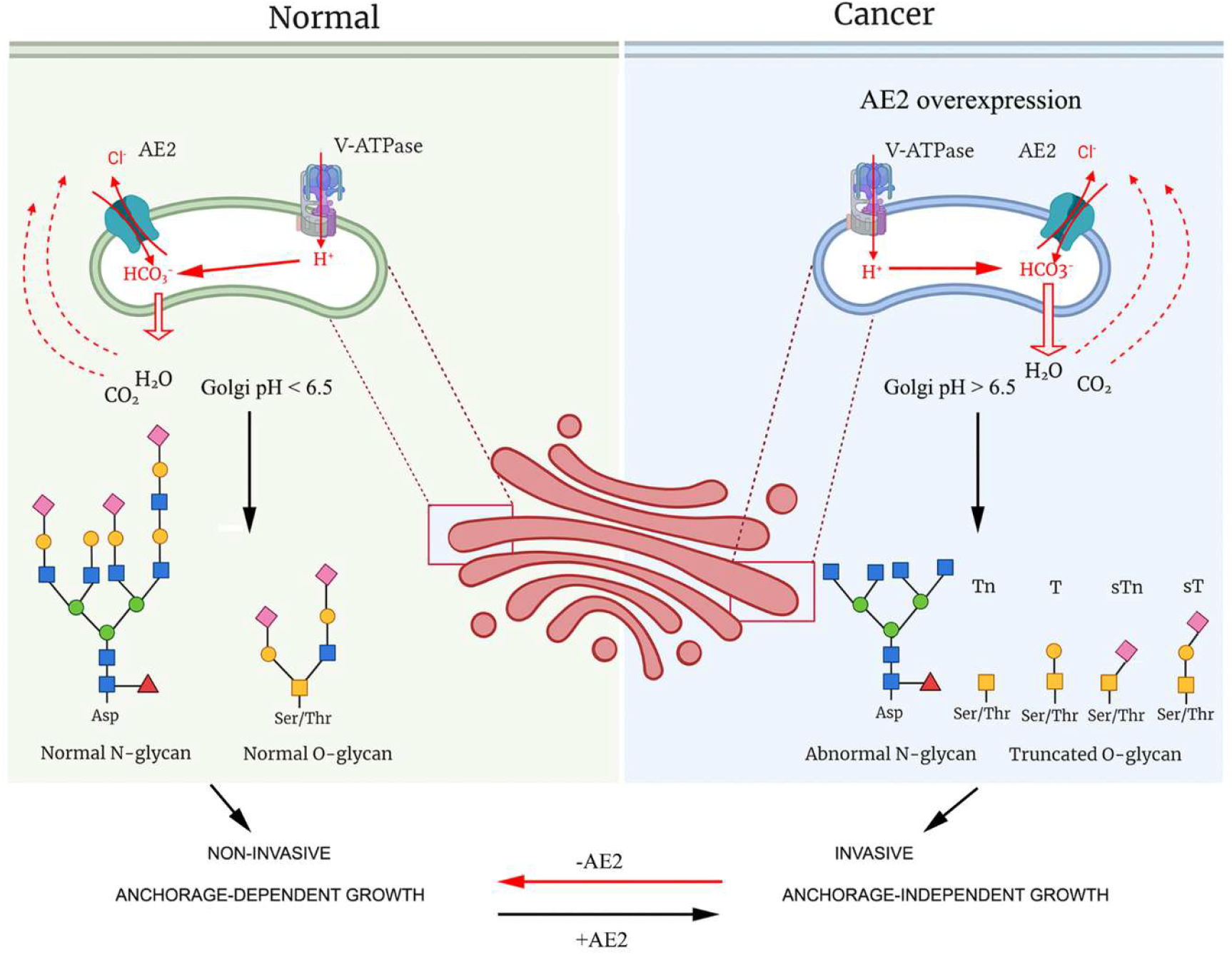
A summary of the AE2-mediated proton leakage pathway and its consequences on cancer cell behavior. For more details see the main text.

## Discussion

We have identified a long-sought proton leakage pathway in the Golgi membranes that is mediated by the AE2a bicarbonate-chloride exchanger. This electroneutral exchanger controls proton leakage rate across Golgi membranes via importing bicarbonate anions in exchange for luminal chloride anions. Strong support for this view was provided by showing that proton leak is strictly dependent on the presence of both bicarbonate and chloride anions as well as on the expression level of the AE2 protein in the cells. Thus, an increase in the concentrations of these anions or AE2 upregulation in the cells induced a concurrent increase in Golgi resting pH and proton leakage rate across Golgi membranes.

Based on these observations, we proposed a model (Fig. 3) in which protons can leak out as water (H_2_O) molecules. Water is produced together with CO_2_ when imported bicarbonate anions spontaneously combine with luminal protons via the well-known bicarbonate-based buffering reaction (see supplementary video S1). It is almost certain that the two end products (H_2_O and CO_2_) then become rapidly expelled across Golgi membranes either via aquaporin channels which can transport, besides water molecules, also other small molecules including glycerol, urea, nutrients, metabolic precursors, waste products, toxins and even gases such as CO_2_ [36,37]. Another option for their egress is diffusion, as the plasma membrane is highly permeable at physiological temperatures to small sized non-ionic water molecules even in the absence of aquaporin channels.

In this respect, it is interesting, and perhaps not even unexpected, to note that Golgi cisternal membranes have a flattened morphology. This shape is optimal for gas exchange as it increases markedly surface-volume ratio, and thereby can speed up egress of the end products from the Golgi lumen. In fact, the need to prevent water and carbon dioxide from accumulating in the Golgi lumen and at the same time, keep the buffering system in balance, may be one of the main reasons for their unique morphology in mammalian cells. In further support of the model, bicarbonate-dependent buffering system is universally utilized for various purposes among living organisms [35]. Moreover, the pKa of the bicarbonate buffering system (pKa 6.1) is within the same range than the resting pH of the medial- and trans-Golgi/TGN (pH 6.5-6.0).

The membrane topology the AE2 anion exchanger and its pH-dependent exchange activity [34] are also consistent with the suggest model. Both the N-and C-termini of the Golgi-localized AE2 face the cytoplasm, i.e. in an identical manner to the plasma membrane-localized AE isoforms. Since all AE2 isoforms function against alkaline loads, they export bicarbonate (a weak base) from the cytoplasm. Therefore, the preferred transport mode of the Golgi AE2 anion exchanger is towards bicarbonate import. The Golgi localized GPHR chloride channel [21] guarantees the availability of luminal chloride anions for bicarbonate import. Considering pH-dependent anion exchange activity of the AE2, previous studies have shown that AE2 exchanger is active above pH 5.0 [43]. Therefore, it retains its activity in the Golgi, the resting pH of which is above pH 6.0. Yet, the pH sensitivity of the AE2 protein may act a safeguard mechanism to prevent other more acidic organelles compartments (such as secretory vesicles or lysosomes) from alkalinizing in a case AE2 might get mislocalized there. These observations as well as its almost exclusive localization only in the Golgi membranes suggests that the AE2-mediated proton leakage pathway applies only to the Golgi compartment and perhaps also to the endoplasmic reticulum (ER), as AE isoforms can acquire their anion exchange activity shortly after their synthesis in the ER [44].

The functional relevance of the AE2-mediated proton leakage pathway was verified by showing that both the AE2 mRNA and protein levels are often upregulated in cancers and in established cancer cell lines. In SW-48 cells, this could be due to the amplification of the chromosome 7 (47+XX, +7), the chromosome AE2 gene is located in humans [45]. Alternatively, it could be caused by epigenetic changes or environmental factors such as hypoxia [12, 46,47]. Nevertheless, upregulation of the AE2 protein was found to have important consequences on cell behaviour most likely via its ability to modulate Golgi resting pH. We showed that high AE2 protein level in SW-48 cells was associated with near neutral Golgi resting pH, and that AE2 knockdown restored it to near normal levels. Moreover, AE2 protein expression level was found to correlate strictly with the Golgi resting pH in COS-7 cells (Fig. 1). These observations provide strong support for the view that upregulated AE2 in certain cancer cells is responsible for their elevated Golgi resting pH. In the other cell types, which display more normal AE2 protein levels, less active V-ATPase or more active CA II could similarly be responsible for their elevated Golgi resting pH. In support of this possibility, immunoblotting data with the CA II showed that this bicarbonate (and proton) producing enzyme is upregulated in MCF-7 cells (Fig. S6).

We also noticed that AE2 knockdown in SW-48 cells rendered cells non-invasive and unable to grow in soft agar (Fig.7). In contrast, the cells proliferated normally when they were grown on normal cell culture plates (Fig. 6). Thus, AE2 knockdown was able to reverse at least two phenomena that are typical for malignant cells, i.e. their invasiveness and anchorage-independent growth phenotype. In this respect, an important issue is how elevated Golgi resting pH can alter cancer cell malignant behaviour to mimic that of non-malignant cells. Due to the multi-faceted nature and the vast number of pathways that contribute to malignant behaviour, identification of the underlying causes remains a challenge. Yet, altered glycosylation may be a common factor as it can affect multiple targets simultaneously. In support of this possibility, our lectin microarray analyses showed significant differences between non-invasive (AE2 KD, COS-7) and invasive (SW-48 Src, SW-48 WT) cancer cells. In non-invasive AE2 knockdown cells the levels of sialylated or non-sialylated T-antigen (a truncated core 1 mucin type O-glycan) were lower than their levels in control cells (SW-48 Scr). More profound changes were detected when wild type SW-48 and COS-7 cells were compared with each other despite the main glycan species in the cells were rather similar between these cells.

Another common feature between non-invasive and invasive cells was the markedly elevated levels of terminally galactosylated complex type N-glycans (Fig. 8). In this respect, it is interesting that most cell surface cell adhesion receptors (such as integrins and selectins), receptor tyrosine kinases, death receptors and matrix metalloproteinases are all glycosylated and that their glycosylation status can modulate their activity as well [46,48,49]. For example, integrins which consist of variable α and β heterodimers, contain multiple potential N-linked glycosylation sites on each subunit. The β1 subunits, which are best characterized, normally carry the same *N*-acetyllactosamine (Galβ(1,4)GlcNAc) type multiantennary structures [49] that was recognized by the RCA 120 lectin and shown to be present at much higher levels in non-invasive AE2 knockdown and COS-7 cells than in invasive cells (SW-48 WT or SW-48 Src). Moreover, changes such as capping the N-acetyllactosamine structure with either α(2,3)-or α(2,6)-linked sialic acid and β(1,6)-GlcNac branching of the N-glycan were shown to have profound effects on cell adhesion and motility [49,50]. Thus, altered N-glycosylation, as revealed with the RCA 120 lectin, may explain in part different behaviour of non-invasive and invasive in a soft agar assay. Another example that supports the role of altered glycans in modulating tumour cell malignant behaviour is the matrix metalloproteinase MMP14, the main enzyme promoting cell invasion into adjacent or distant tissues. Recently, Bard and his group [48] showed that in mouse liver cancer model, O-glycosylation of MMP14 by the initial GalNAc residue is markedly enhanced due to relocalization of relevant GALNT glycosyltransferase to the ER. This increase in turn markedly increased MMP14 activity as well as tumour cell invasion and tumour expansion. A third supporting example is TRAIL induced apoptosis, as it is also regulated by altered glycosylation [46]. In this case, impaired O-glycosylation and in particular, extension of the (sialyl)Tn-antigen to (sialyl)-T-antigen (with added galactose residue) of death receptors was found to confer apoptosis resistance to cells via affecting receptor oligomerization state. This consistent with our finding that AE2 knockdown cells showed fragmented nuclei when grown in soft agar.

Consistent in part with this, we observed a significant decrease in the Tn- and T-antigen levels (as well as of a-linked GalNAc) in non-invasive COS-7 cells compared to invasive SW-48 WT cells with ACA, PNA, SNA-II, PTL-I and DBA lectins (Fig. 8D). The only exception was the VVL lectin (specific for GalNAc), which showed an increase. In AE2 knockdown cells, however, we did not detect a similar decrease with these same lectins, except with the ACA lectin that is specific for the T-antigen (i.e. Gal(β1,3) GalNAc; Fig. 8C). Thus, based on glycan profiling, the T-antigen, rather than the Tn-antigen, appears to be more critical determinant for cancer cell invasion. Nevertheless, further studies are needed to clarify this issue with different cell lines and/or by the use of mass spectrometry to allow more detailed glycan analyses. This might be challenging, as defining critical glycans for invasion may require in the worst case both protein- and glycosylation site-specific glycan analyses [50].

In summary, we find that overexpression of the AE2 anion exchanger in certain cancer cells is responsible for their elevated Golgi resting pH, a phenomenon that was shown to impair glycosylation events in the Golgi. Altered glycosylation in turn promotes cancer cell invasiveness and anchorage-independent growth and perhaps even anoikis-resistance, as it is also regulated by O-glycosylation [46]. How glycosylation changes modulate these events remain unclear, but they seem to be important for the activity and/or functioning of various cell surface constituents that regulate cell adhesion, motility, invasion and colonization into adjacent or distant organs. This view does not contradict the importance of cancer-associated mutations in tumorigenesis, given that they are necessary for the cell proliferation and tumour growth, while cell surface glycans seem to have their main roles in promoting tumour progression, i.e. when benign cells become malignant. Our findings also emphasize that tumour cell malignant behaviour can be reversed by normalizing Golgi resting pH and its pH-dependent glycosylation potential. In this respect, our findings may open new possibilities in the future to prevent metastatic spread of cancers, the main cause of death of cancer patients. In the worst case, they can provide new diagnostic tools for detecting metastasis-prone tumours and thereby help patient care.

## Materials and methods

### Reagents, antibodies and plasmids

All reagents were purchased from Sigma-Aldrich (St. Louis, MO, USA) unless stated otherwise. The antibodies against the Golgi marker GM130 was purchased from BD Biosciences (#610822, San Jose, CA, USA). The horseradish peroxidase-conjugated secondary antibodies were from Abliance (Compiègne, France). An antibody against AE2 C-terminal peptide was prepared and affinity-purified as described earlier [51]. Alexa Fluor488- and 594-conjugated anti-mouse and anti-rabbit secondary antibodies and AlexaFluor488-or Alexa594-conjugated peanut agglutinin (PNA) were purchased from Molecular Probes (Eugene, OR, USA) or from Invitrogen (Carlsbad, CA, USA). Unconjugated lectins were purchased from either EY laboratories (San Mateo, CA, USA) or Vector Laboratories Inc. (Burlingame, CA, USA). Anti α-tubulin antibody was from Sigma Aldrich (St. Louis, MO, USA)

The plasmid encoding the medial/trans-Golgi localized pHluorin [33] was a kind gift from Dr. G. Miesenböck (Oxford, UK). AE2-mCherry plasmid was prepared by sub-cloning the full length AE2 in frame into the pcDNA3-monomeric Cherry (mCherry) vector (Invitrogen, Carlsbad, CA, USA).

### Cell culture and transfections

All cell lines (COS-7, HeLa, HT-29, SW-48, CaCo-2, DLD-1, MCF-7, MDA-MB 231, RCC4, A431) were from ATCC (Manassas, VA). Cells were grown in high glucose Dulbecco’s modified Eagle’s medium (DMEM) supplemented with Glutamax (Gibco BRL, Grand Island, NY, USA), 10% fetal bovine serum (HyClone, Cramlington, United Kingdom) and antibiotics (100 U/ml Penicillin and 100 μg/ml Streptomycin; Sigma-Aldrich, St. Louis, MO) in humidified conditions at +37°C and at 5% CO_2_. Transfections were performed using either the Lipofectamine^®^ 3000 (Thermo fisher scientific, CA, USA) or the FuGENE 6™ (Promega, Fitchburg, WI, USA) transfection reagents according to the suppliers’ instructions for 1-2 days before the experiments. In certain cases, cells were also transfected with electroporation using Amaxa™ Nucleofector Kit R (Cat.no.VCA-100) and the Nucleofector™II device (program W-001, Lonza Group AG, Cologne, Germany) as suggested by the manufacturer.

### Immunofluorescence electron microscopy

Immunofluorescence microscopy was performed by fixing cells with 2% p-formaldehyde (30 min), after which cells were permeabilized with 0.1% saponin in PBS and stained with the anti-GM130 antibody or with AlexaFluor488-or Alexa594-conjugated lectins (1h at RT). After washing, cells were treated with relevant species-specific Alexa Fluor488- and 594-conjugated anti-mouse or anti-rabbit secondary antibodies. After mounting, cells were imaged using Olympus BX 51 microscope with appropriate filter sets for the dyes. Alternatively, Zeiss Observer Z1 confocal microscope (LSM 700, Carl Zeiss AG, Oberkochen, Germany) with appropriate filter sets, the Zen2009 software, a 63X Plan-Apo oil-immersion objective, was used. Co-localization studies of the mCherry fusion constructs or GT-pHluorin constructs with the Golgi marker (GM130) antibody were done using Alexa488 or Alexa 594-conjugated secondary antibodies, respectively.

### Immunoblotting

Immunoblotting of the α-tubulin (used as a loading control) were carried out by using sodium dodecyl sulphate polyacrylamide gel electrophoresis (SDS-PAGE). In brief, cells were lysed on plates with the lysis buffer (50 mM Tris-HCl (pH 7.5), 150 mM NaCl; 1% TX-100; 2 mM EDTA; 2 mM EGTA) supplemented with protease inhibitors (Complete Mini, Roche, Basel, Switzerland). After clearing (15000×g for 15 min), 50-100μg of total protein in SDS-sample buffer was separated using 7.5% SDS-PAGE gels.

Blue-native polyacrylamide gel electrophoresis was used for immunoblotting of the AE2a protein as described below. Cells were washed twice with PBS, scraped from the dishes, pelleted by centrifugation and lysed on ice in TND/TX-100 buffer (25 mM Tris pH 7.5, 150 mM NaCl, 0.5% Deoxycholic acid, 1% Triton X-100) for 1 h. After clearing by centrifugation (15.000g, +4C, 15 min), the TX-100 soluble fractions were mixed with 5x Tris/Glycerol native sample buffer before loading onto a 4%-15% Mini-Protean TGX gel (Bio-Rad, Hercules, CA). The samples were run in Tris/Glycine buffer supplemented with 0.02% Coomassie Brilliant BlueG-250 until the samples had migrated 1-2 cm into the gel, and thereafter without the dye at 20 mA for 2 h. The samples were transferred onto the PVDF membrane (Bio-Rad) with Bio-Rad Trans-blot Turbo for mixed molecular weight program (1.3A-25V; 7 min). The membrane was quenched by using 5% non-fat milk in TBS-Tween (50 mM Tris-150 mM NaCl-0.02% Tween 20, pH 7.6) supplemented with 0.1% BSA overnight at +4C. The blot was then incubated with a polyclonal anti-AE2 C-terminal antibody (anti-AE2Ct) or the mouse monoclonal anti α-tubulin antibody (Sigma Aldrich, St. Louis, MO, USA) in TBS-Tween 20 + 0.1% BSA) for 2 h at RT. After washing 3 times for 10 min with TBS-Tween 20, rabbit anti-mouse or goat anti-rabbit HRP-conjugated secondary antibodies (Abliance, Compiègne, France) were added in TBS-Tween-0.1% BSA for 1 h at RT (1:10.000 dilution). After final washings (4 times 15 min in TBS-Tween), protein bands were visualized using the ECL reagent and the GelDoc instrument (both from Bio-Rad, Hercules, USA) before quantification using the build-in Image Lab-software.

### Knockdown of AE2a in COS-7 and SW-48 cells

Given the lack of hereditary human diseases related to human AE2, and the severe phenotype described for SLC4A2 (AE2) knockout mice [27], we decided to utilize an inducible shRNA-mediated gene silencing of the AE2a protein (as well as other AE2 transcripts) that also allowed us to control AE2 expression level in the cells. Cells were transfected with the AE2-specific and control (scrambled) shRNAs using the inducible SMART-vector constructs (Dharmacon, Lafayette, CO, USA) as suggested by the manufacturer. The shRNA sequences are shown in Table S1. Stably transfected cells were selected first against puromycin (Gibco, 0.75-1.5 μg/ml, 48 h) before repeated sorting of cells by fluorescent activated cell sorter (BD FACSAria™) and collecting cells with high (>10-fold over the background) doxycycline-inducible RFP expression. Cell lysates were then prepared from the stable transfectants with or without doxycycline induction (50-250 ng/ml) for 72 h and used for the determination of the AE2a protein expression levels by immunoblotting of the cell lysates with the AE2 C-terminal antibody as describe above.

### Lectin microarray analyses

Cells cultivated on plates to 70-80% confluency were lysed for 30 min on ice with lysis buffer (50mM sodium tetraborate buffer, pH 8.5, 150mM NaCl, 1% Triton X-100) supplemented with the protease inhibitor cocktail (Complete Mini, Roche) and clarified by centrifugation (12,000×*g* at 4 °C for 15 min). 12 μg of total protein of each cell lysate was labelled with 6 μg of NHS activated DyLight 630 dye (Thermo Scientific, Waltham, USA) in 50 μl of labelling buffer (50 mM boric acid/ 150 mM NaCl, pH 8.5) for 1h at RT with constant agitation (600 rpm). The reaction was quenched at RT for 1 h by adding 50 μl of quenching buffer (75 mM ethanolamine in 200 mM Tris-HCl/150 mM NaCl, pH 8.5) before dilution (1:12) with the assay buffer (50mM Tris/300mM NaCl/2mM MgCl_2_/2mM MnCl_2_/2mM CaCl_2_, pH 7.1) to give the final pH 7.4 to the labelled sample mix. After clearing the mix by centrifugation (12,000×*g* for 10 min at RT), 400 μl of the labelled sample was applied into each well on pre-printed and pre-quenched (with 50 mM ethanolamine) Nexterion H microarray slides (Schott, Germany). The labelling was further incubated in a humidified chamber with constant agitation for 2 h in RT. Slides were then washed 5 times for 5 min each with the washing buffer (50 mM phosphate buffer/0.05% Tween). Array images was generated using the Genepix 4200AL laser scanner (Axon Instruments) using an appropriate filter set for the DyLight 633™ dye. The mean intensities of bound label were quantified in triplicate from 4 parallel arrays (36 measurement spots/sample) using the GenePix Pro® microarray analysis software. Sugar binding specificities of the lectins were deduced from the manufacturer’s product sheets. The corresponding lectin names are available upon request. The specificity of the lectins for their own glycotops was verified during optimization of the protocol by using labelled fetuin and asialofetuin as markers.

### Golgi pH measurements

Golgi resting pH in the cells was determined as follows. Briefly, after equilibration in the assay buffer (PBS supplemented with 0.5 mM MgCl_2_, 0.9 mM CaCl_2_ and 4.5 g/l D-Glucose (PBS/D-Glucose) at RT without CO_2_, the Golgi region of the cells expressing the GT-pHluorin were imaged by using the Operetta™ high content imaging system (PerkinElmer Inc.). I*n situ* pH calibration was done by replacing the bath solution with different pH calibration buffers (125 mM KCl, 20 mM NaCl, 1.0 mM CaCl_2_, 1.0 mM MgCl_2_) pre-adjusted to pH 7.5, pH 6.5 and pH 5.5 with 20 mM HEPES, MOPS or MES, respectively. Ionophores (5 μM nigericin, 5 μM monensin) were included in the buffers to dissipate the existing monovalent (H^+^, K^+^, Na^+^) ion gradients.

Golgi acidification and leakage rate measurements were performed in triplicate as follows. Cells expressing the pHluorin were equilibrated as above, imaged and then permeabilized with Streptolysin O (SLO, 3 μg/ml) either in chloride-free or chloride-containing high potassium buffer (120 mM KCl, 30 mM NaCl, 10 mM EGTA, 10 mM MgCl_2_, 10 mM Hepes, pH 7.2). Chloride-free high potassium buffer contained corresponding gluconate/sulphate salts instead of chloride anions, which cannot be used for AE2a-mediated bicarbonate exchange [43]. Bicarbonate was added in the chloride-containing bath medium just before the assay was started. Golgi acidification rate (a sum of proton pumping and its leakage rates) was followed adding fresh ATP (10 mM) to the bath solution. Ratio imaging (15 sec intervals) of the selected Golgi regions was performed until the ratios (Golgi pH) reached a plateau. The Zeiss Axio Observer Z1 microscope equipped with a dual FRET camera system and appropriate filter sets for two different excitation (420 and 470 nm) and one emission (500– 550 nm) wavelengths was used for imaging. Proton leakage rates were then measured by replacing the ATP-containing buffer with an ATP-free buffer with added concanamycin A (CMA, 1 μM), a potent V-ATPase inhibitor. Proton leakage (i.e. Golgi pH increase) was then followed again by ratio imaging until it reached a plateau. At the end of each experiment *in situ*-pH calibration was performed by using pre-calibrated calibration buffers (pH 5.0, 5.5, 6.0, 6.5, 7.0 and 7.5) as above. Intensity ratios were then transformed to pH values using the determined formulas for the obtained sigmoidal calibration curves. Numerical data were processed using Microsoft® Excel solver (Redmond, WA, USA).

### Immunoblotting of human colorectal cancer tissue samples

Human cancer tissue specimens were obtained from the Pathology research unit, Oulu University Hospital following Institutional Ethical Committee Review Board guidelines (permission numbers. 25/2002, 42/2005, 122/2009). Fresh samples were frozen and stored at −80°C until use. Proteins were extracted using 500 μl of ice cold lysis buffer (50mM Tris-HCl, pH 7.4, 150 mM NaCl, 1% Triton X-100, 0.5% deoxycholic acid with a protease inhibitor) and pre-cooled tissue homogenizer (TissueLyser LT, Qiagen GmbH, Hilden, Germany) for 2-4 min at 45 Hz. After homogenization, the lysates were clarified by centrifugation (15 000 rpm, 15 min at +4°C) and run on a Blue-native gel electrophoresis (for AE2) and SDS–PAGE (for α-tubulin) as described in the main text.

### Cell proliferation and wound healing migration assays

The assays were performed using the IncuCyte® Live-Cell Analysis System and established protocols (Essen BioScience, Newark Close, and UK). In brief, for cell proliferation assays, wild type SW-48 cells and cells stably transfected with the scrambled or AE2-specific shRNAs plasmids were plated into special 96-well plates (5 × 10^3^ cells/well), and cultured in DMEM supplemented with 10% FCS and penicillin-streptomycin for six days. Cells were imaged at 2 h intervals during the 5-day culture period before quantification with the IncuCyte® software.

For the scratch wound cell migration assay, 5 × 10^4^ cells/well were plated and after reaching confluence, cell monolayers were grown on Image Lock 96-well microplates (Essen Bioscience) in DMEM with 1% FCS after scratching 700-800 μm wide “wounds” using the 96 pin IncuCyte® Wound Maker (Essen Bioscience). Wound closure (migration) was then followed by using phase contrast imaging for 6 days at 2 h intervals. Basic IncuCyte® software and settings were used for the analyses. The data are expressed as percentages of confluence in each well.

### 3D invasion assay

The invasive properties of wild type SW-48 and AE2 knockdown cells as well as COS-7 cells stably overexpressing the AE2a variant (G418 selection) were investigated using an stablished organotypic 3D-myoma-invasion model [39]. In brief, myoma discs pre-equilibrated at +4°C in DMEM, were placed in tightly fitted Transwell® inserts (Corning, Inc., Corning, NY, USA) after which 5 × 10^5^ cells (in 50 μl of DMEM) were added on top of each disc. After attachment, myoma discs with cells were transferred onto uncoated nylon discs placed on curved steel grids (3 × 12 × 15 mm) in 12-well plates, each well containing 1ml of fresh media with and without doxycycline. The myoma organotypic cultures were maintained for 14 days with daily media changes. Each assay was performed in triplicate. The specimens were fixed in 4% formalin overnight, dehydrated, and embedded in paraffin. Finally, 6 μm thick sections were cut and deparaffinized before staining with Mayer’s Hematoxylin-Eosin. After imaging, the invasion depth of all invaded cells (area) in each microscopic field was determined by measuring the distance of the cell invasion front from the top cell layer on each disc using the Image J (Fiji) v1.46o (National Institute of Health, USA).

### Soft agar colony formation assay

Black wall CellCarrier-96 Ultra Microplates (PerkinElemer, Inc, Waltham, Ma, US) were used in the soft agar colony formation assay. Prewarmed 25 μl of 2 × Dulbecco’s modified Eagle’s medium (D-MEM; containing 20% FBS, 200 U/ml penicillin, 200 μg/ml streptomycin and 25 μl of melted 1% Ultra Pure™ agarose (Thermo fisher scientific, CA, USA) solutions were mixed and transferred into each well and put at 4 °C for 30 min to allow the agar layer to solidify. Trypsinized cells (2.5x 10^3^ cells) were then suspended in 25 μl of D-MEM/10% FBS and mixed with 25 μl of 2 × D-MEM containing 20% FBS and 25 μl of 0.7% agar before adding the suspension on top of the solidified agar bed, and cooling at 4 °C for 15 min. After adding 50 μl of 1 x DMEM per well, the plates were incubated with for 30 days at 37 °C and 5% CO_2_. The medium supplemented (or not) with 100 ng/ml doxycycline was changed every 2-3 days. Thirty days post-seeding, cells were fixed with 2% PFA at RT for 30 min and stained with the Hoechst 33342 dye (1 μg/ml) at 37 °C and 5% CO_2_ before imaging with the Operetta™ high content imaging system (PerkinElmer Inc.). Forty fields were acquired from each well using appropriate filter sets, and a 20× objective. Images were analyzed using the Harmony software with selected scripts to allow segmentation to cell colonies. Segmentation criteria included on colony size (>3 cells/colony), circularity and fluorescence intensity of nuclei.

### Statistical analyses

Statistical analyses between control and test groups were performed using GraphPad Prism Software (GraphPad Software Inc., La Jolla, CA, USA) and the 2-tailed Student’s *t*-test. Probability values smaller than p < 0.05 (*) were considered significant.

## Abbreviations

GlcNAc: N-acetylglucosamine
GalNAc: N-acetylgalactosamine
Gal: Galactose
Man: Mannose
Fuc: Fucose

## Data availability

The source data files and computer codes used in this manuscript are available from the corresponding author upon reasonable request.

## Acknowledgements

The authors wish to thank Dr. Antti Hassinen for his help in setting up the protocols for Operetta High Content Imaging System. We also acknowledge Maija-Leena Lehtonen and Tanja Kuusisto for expert technical assistance, M.Sc. Teemu Viinikangas for setting up the Genepix 4200AL laser scanner protocols. We also greatly acknowledge the University of Oulu Graduate School, Finnish National Agency for Education (CIMO), the Thelma Mäkikyrö Foundation and the Academy of Finland for funding.

## Author contributions

EK and AR performed the experiments, MR and TS provided help in myoma invasion assays. AT and MM provided the human colorectal tissue specimens with low-grade histology. Their use has been accepted by the Oulu University Hospital ethical committee (License numbers: 25/2002, 42/2005, 122/2009). EK and SK designed the experiments, analysed the data, made the figures and wrote the manuscript.

## Additional information

Video S1 (see a separate file)

