## Supplementary material for "SLC4A2 Anion Exchanger Promotes Tumor Cell Malignancy via Enhancing H^+^ Leak across Golgi Membranes": supplemetary materials

**Fig. S1.** **Distribution of the AE2a protein in the Golgi sub-compartment in COS-7 cells**

**Fig. S2 legend. The effect of pH and bicarbonate buffers on Golgi resting pH**

**Fig. S3. The effect of Golgi fragmentation, hypoxia on Golgi resting pH**

**Fig. S4**. **AE2 Knockdown efficiency in SW-48 cells**

**Fig. S5. Lectin microarray glycan profiles of non-invasive cells and invasive cancer cells**

**Fig. S6. Expression level of carbonic anhydrase II (CAII) in different cancer cell types**

**Table S1.** **Sequences and location of the AE2-specific shRNAs used for AE2 knockdown with doxycycline-inducible SMART-vector system**.

**References for supplementary information**

**Fig. S1.**

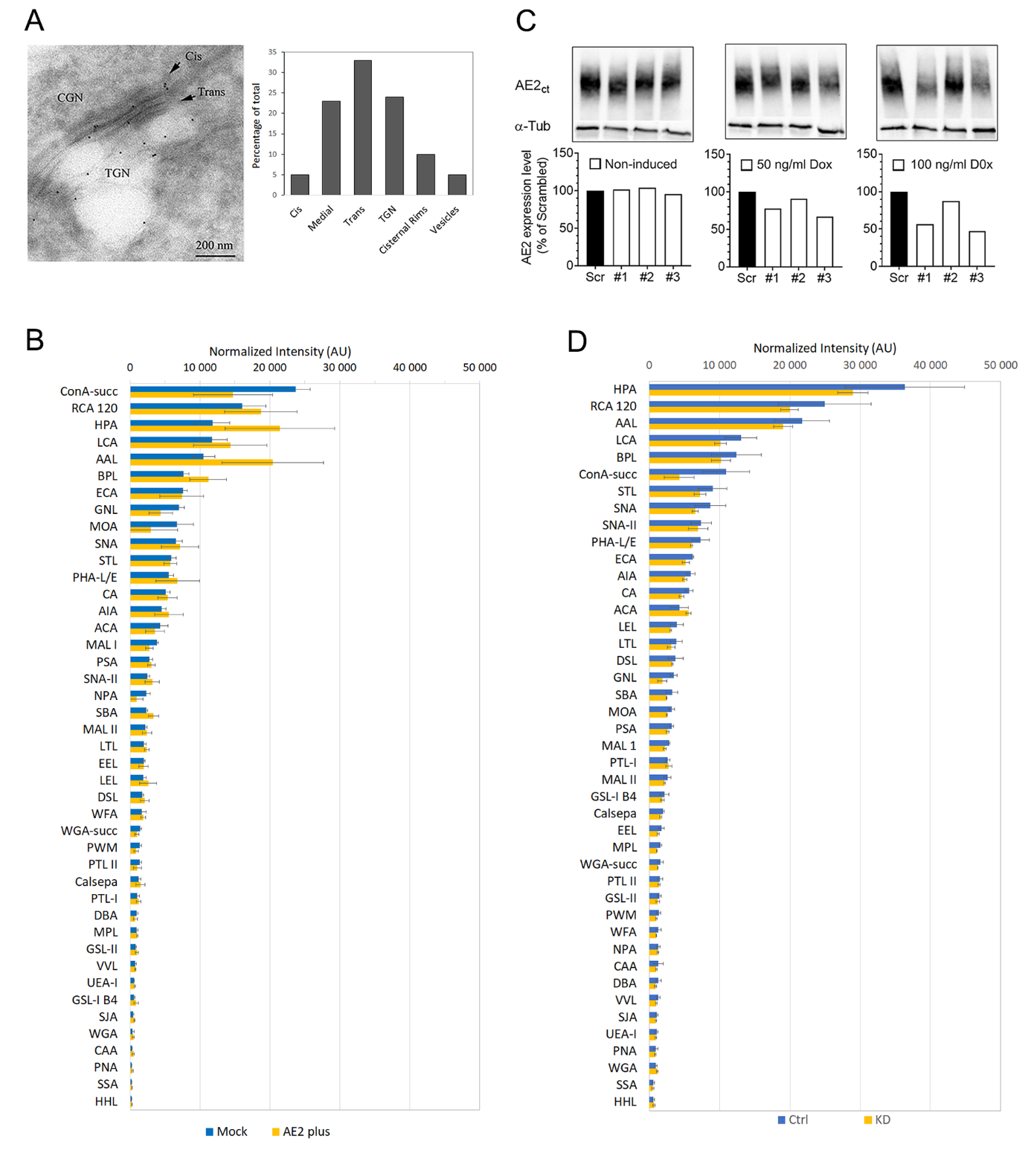

**Fig. S1.** **Distribution of the AE2a protein in the various Golgi sub-compartment in COS-7 cells.** **(A)** Distribution of the AE2a protein in the Golgi sub-compartments was assessed by electron microscopy and immune-gold staining. A representative figure is shown. In brief, cells were processed according to the established protocols for cryoelectron microscopy. Thin cryo-sections were cut and incubated with the anti-AE2Ct antibody and then with 10 nm diameter gold particles coated with protein A. Sections were examined using the Philips CM100 transmission electron microscope. (Right) Quantification of the gold particles in each Golgi sub-compartment was done according to Rabouille et al. [1]. Altogether, 984 particles were used for the quantification). **(B)** Lectin microarray glycan profiles of AE2a-mCherry overexpressing cells. Mock-transfected cells were used as a control. Total binding intensities are shown as the means (± S.D.) after sorting from largest to smallest and normalization of the data values against α-tubulin. **(C)** Knockdown of the AE2a protein in COS-7 cells with different AE2 shRNAs and with a control plasmid (scrambled shRNA). Doxycycline at 50-100 ng/ml was used for testing their potency. Immunoblotting of cell lysates prepared from either non-induced or induced COS-7 cells stably expressing the AE2-specific or scrambled shRNAs. For the induction, 50-250 ng/ml of doxycycline was used for 72 h before preparing cell lysates. ImageJ was used for quantification of the AE2a protein and α-tubulin in the blots. The intensity values were normalized against α-tubulin levels. **(D)** Lectin microarray glycan profiles of AE2a knockout (shRNA#1) and control cells that were transfected with the scrambled shRNA. Total binding intensities are shown as the means (± S.D.) after sorting from largest to smallest and normalization of the data values against α-tubulin.

**Fig. S2**

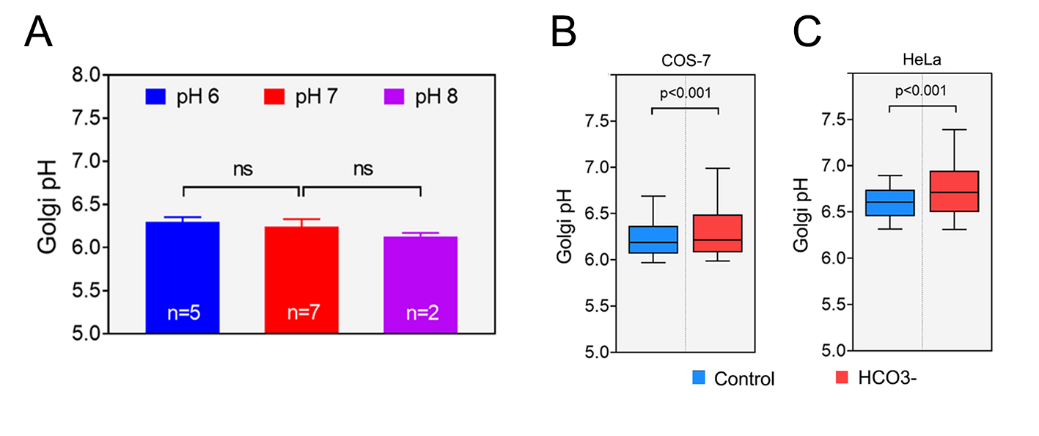

**Fig. S2 legend. The effect of pH and bicarbonate buffers on Golgi resting pH. (A)** Cells expressing the GT-pHluorin were equilibrated for 20 min in different media pre-adjusted to pH 6, 7 or 8 using MES, MOPS and Hepes, respectively. Cells were then ratio-imaged to determine their Golgi resting pH using the exponential standard curve equation as described in the “Materials and methods”-section. The bars represent the means (±SD) of separate experiments depicted in the graph. The mark “ns” denotes to p-values > 0.05. (**B-C**) The effect of bicarbonate (20 mM) on Golgi resting pH in intact COS-7 cells (B) and HeLa cells (C) is shown as a box plot. The whiskers represent 10^th^ and 90^th^ percentiles. In brief, cells were grown either in the absence (blue boxes) or presence (red boxes) of extra bicarbonate (20 mM) for 24 h before determination of Golgi resting pH using the Operetta^TM^ high content imaging system.

**Fig. S3**

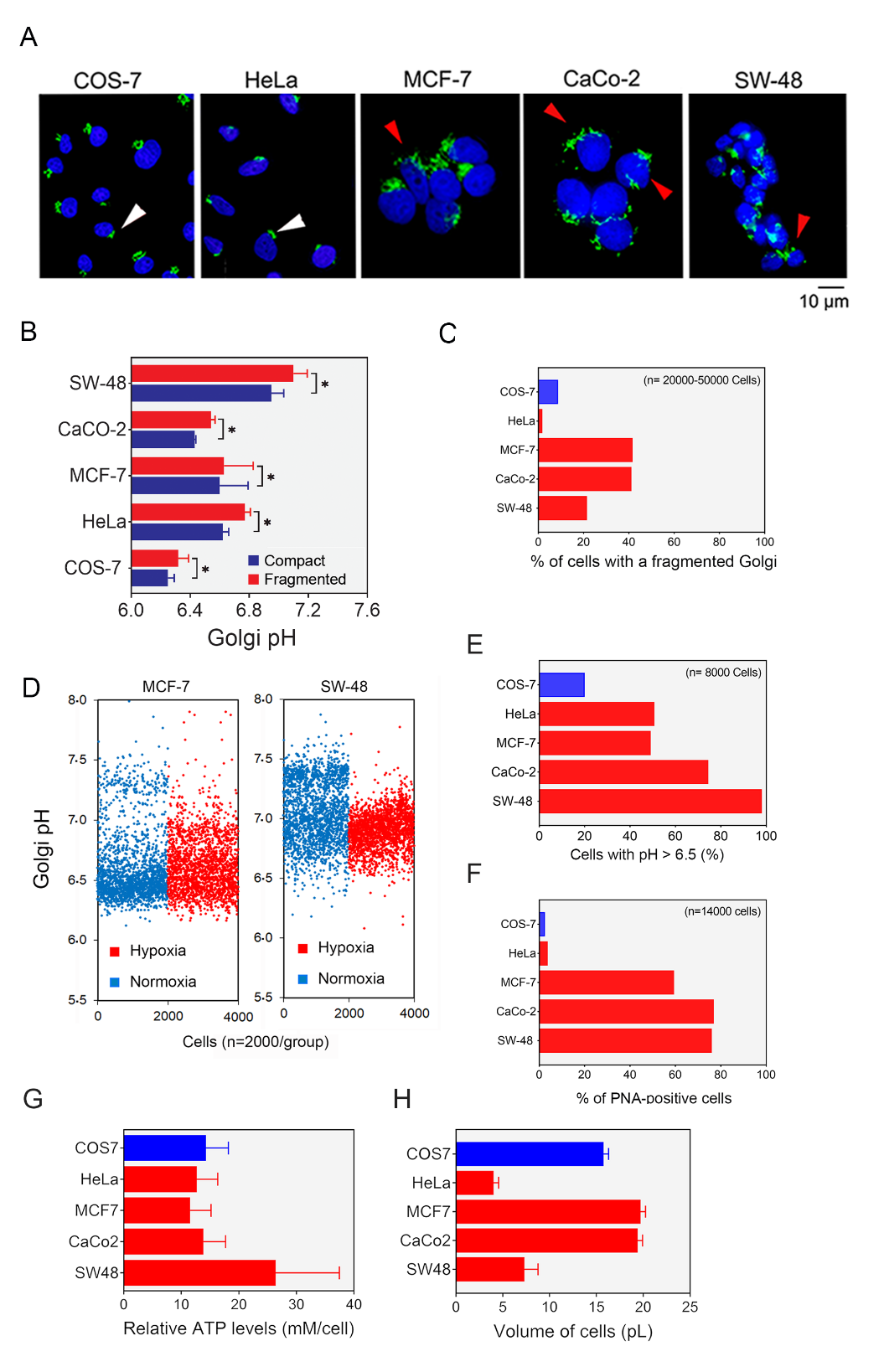

**Fig. S3. The effect of Golgi fragmentation, hypoxia on Golgi resting pH (A-D).** (**A**) Immunofluorescence microscopy analysis showing examples of the Golgi with compact (white arrowheads) or fragmented (red arrowheads) morphology. Golgi elements were classified as fragmented if the number of distinct Golgi elements was equal or higher than two in a single cell (depicted by the nucleus, blue colour). **(B)** Golgi resting pH in compact and fragmented Golgi elements. pH measurements were done as described in the “Materials and Methods” section after selecting cells having either a compact or fragmented Golgi elements. The Harmony software was used for selection. **(C)** Quantification of the cells with the fragmented Golgi morphology in different cancer cell types. Cells having a fragmented Golgi morphology were quantified in each cell population using the Harmony software. The results are presented as percentages of all cells (n= 2-5 x 10^4^ cells) in each cell population. **(D)** The effect of hypoxia on Golgi resting pH in MCF-7 and SW-48 cells. Cells were grown and analysed at hypoxic conditions. The plot shows single cell data values measured with the Operetta high content imaging system. **(E-F) The proportion of cells that display abnormally high Golgi resting pH or PNA binding**. **(E)** The percentage of cells with abnormally high Golgi resting pH in the five cancer cell types tested. Cells were grown on Cell Carrier Ultra 96-well plates, transfected with the GT-pHluorin plasmid, and used for Golgi resting pH 16-20 hr later. For classification, The Golgi resting pH above pH 6.5 was used as a set point. Total cell numbers in either class was counted using the Harmony software. (**F**) The percentage of cells that stain positively with PNA lectin. Cells were grown, fixed and stained with Alexa-Fluor 594 conjugated PNA -lectin and with the Hoechst 33342 dye (to stain nuclei of each cell). For quantification of PNA-positive cells, we selected a signal cut-off of 500 AU units/cell, since the background signal (determined by using unstained cells) was equal or less than 300 AU units/cell. Cell numbers were then counted to get their percentages in each cell population. **(G-H)** **Cellular ATP levels of different cancer cell lines.** **(G)** ATP levels were determined according to the manufacturer's instructions, using the luminescent ATP Detection Assay Kit (Abcam, Cambridge, MA, USA). In brief, cells were grown in white, clear bottom wells in a 96-well plate for 24 h. Thereafter, 50 μl of lysis buffer was added to each well and incubated for 5 min in an orbital shaker at 700 rpm. Then, 50 μl of substrate solution were added to each well and incubated for 5 min in an orbital shaker at 700 rpm. The luminescent signal was measured using the TECAN Infinite 1000 Pro multimode microplate reader (Männedorf, Switzerland). The results are expressed as mM ATP per cell (±SD, n=4) after normalizing the values with the mean cell volume and cell number/well. Operetta high content imaging system was used for cell counting. **(H)** Determination of the cell volumes using fluorescent activated cell sorting (FACS). The raw data values (cell size, µm) was transformed to cell volumes using the equation for a round 3D subject.

**Fig. S4**

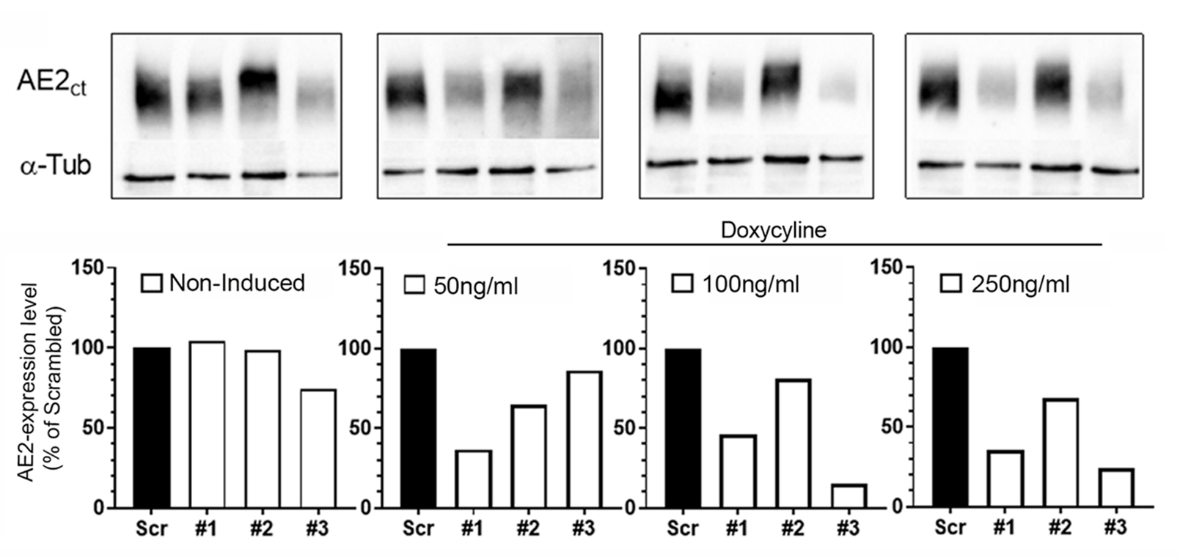

**Fig. S4**. **AE2 Knockdown efficiency in SW-48 cells**. SW-48 cells carrying each one of the AE2-specific shRNA constructs (#1, #2, #3) were used to define the doxycycline concentration that most efficiently attenuates AE2 protein expression. The expression levels were determined by immunoblotting with the anti-AE2 C-terminal antibody either with or without the 3-day- induction using the depicted concentrations of the antibiotic.

**Fig. S5**

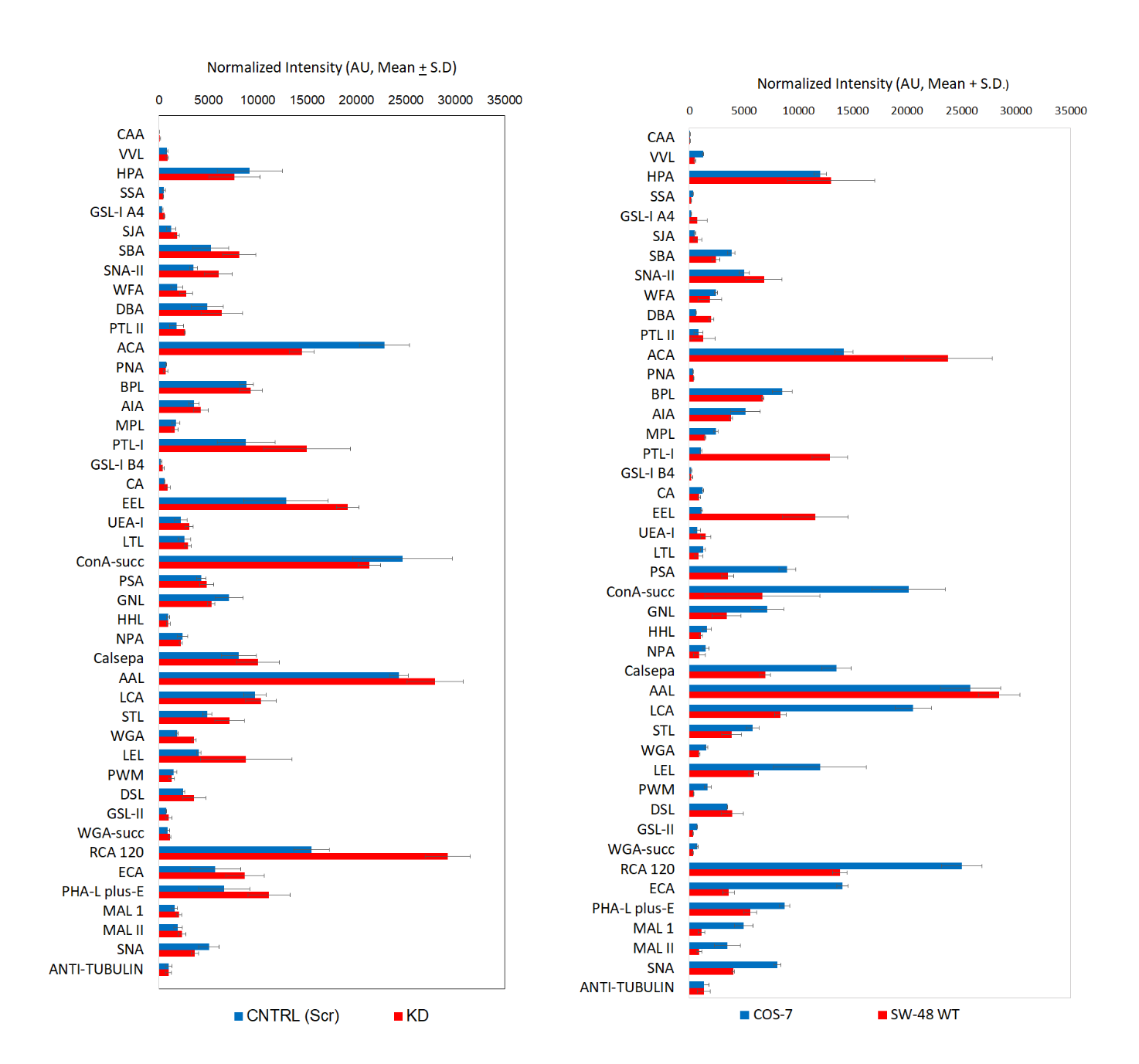

**Fig. S5. Lectin microarray glycan profiles of non-invasive cells and invasive cancer cells.** Total bindings intensities are shown as the means (± S.D.) after normalization against α-tubulin. AE2 knockdown cells (KD) was compared to its own control cells (SW-48 Scr). The glycan profiles of the wild type cells (COS-7, no-invasive) and SW-48 cells (invasive) were also compared with each other.

**Fig. S6**

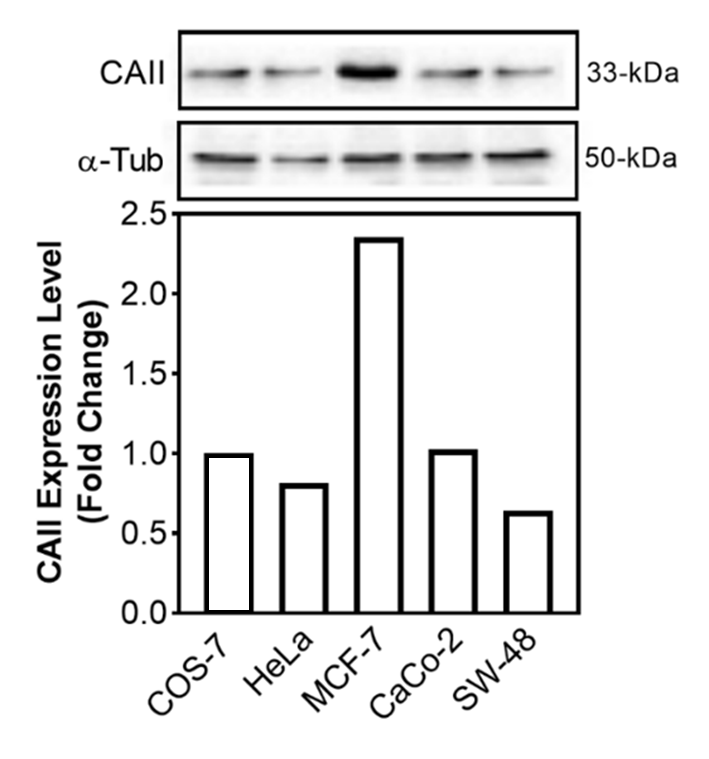

**Fig. S6. Expression level of carbonic anhydrase II (CAII) in different cancer cell types.** Cells were solubilized with SDS-sample buffer, run on a 4-20% gradient polyacrylamide gel and immunoblotted with anti-CAII and anti-α-tubulin antibodies. Quantification of the band intensities was done using ImageJ. The graph shows the normalized CAII levels as fold changes in comparison to that of non-malignant COS-7 cells.

**Supplementary tables**

**Table S1.** **Sequences and location of the AE2-specific shRNAs used for AE2 knockdown with doxycycline-inducible SMART-vector system**. Sequence identity between the five human SLC4A2 transcript variants (1, 2, 3, 4 and X1) that are transcribed from the SLC4A2 gene is 100%. The shRNA sequences locate in exons 12 or 15 in the SLC4A2 gene, corresponding to part of the cytosolic domain of the AE2 proteins. The shRNAs target the following SLC4A2 transcripts: NM_001199692, NM_001199693, NM_001199694, NM_003040, and XM_006716094. No potential off-targets were recognized based on mis-match checking by using Blast-n and the In-Silico Solutions Bioinformatic Tools Site <http://projects.insilico.us/SpliceCenter/Credits.jsp> [2].

| Name | Antisense Sequence | Promoter | Reporter | Chr | Strand | Exon |
| --- | --- | --- | --- | --- | --- | --- |
| shRNA#1 | TCGGCCTTGATCTGGTCAG | mCMV | TurboRFP | 7 | + | 12 |
| shRNA#2 | CCGATCTCGTGGTAGTCCA | mCMV | TurboRFP | 7 | + | 15 |
| shRNA#3 | ACTGCGTCCAACTCCACAG | mCMV | TurboRFP | 7 | + | 15 |
